# Nanoparticle-mediated microRNA-145 Delivery for Vascular Smooth Muscle Cell Phenotype Modulation and Atherosclerosis Treatment

**DOI:** 10.1101/2020.09.09.290361

**Authors:** Deborah D. Chin, Christopher Poon, Jonathan Wang, Johan Joo, Victor Ong, Zhangjingyi Jiang, Kayley Cheng, Anastasia Plotkin, Gregory A. Magee, Eun Ji Chung

## Abstract

Vascular smooth muscle cells (VSMCs) change from contractile to the synthetic phenotype during atherogenesis and 30-70% of cells that make up plaques have been elucidated to be of VSMC origin. MicroRNA-145 (miR-145) is responsible for regulating VSMC phenotypic switching, and low miR-145 levels in circulation have been linked with atherosclerosis. Hence, we developed nanoparticles for targeted delivery of miR-145 by synthesizing micelles co-assembled with miR-145 and the CCR2-binding peptides for plaque targeting. The miR cargo was protected in micelles from premature endosomal degradation and rescued contractile markers in synthetic VSMCs and SMCs isolated from patient arteries *in vitro*. In ApoE-/- mid-stage atherosclerotic mice, miR-145 micelles halted plaque growth and maintained contractile phenotypes similar to baseline levels. In early-stage atherosclerosis, a single dose of miR-145 micelles prevented lesion growth by 49%. We present the potential of miR-145 micelles as a therapeutic that can be applied longitudinally and intervene throughout atherosclerosis pathogenesis.

## Introduction

Vascular smooth muscle cells (VSMC) are the predominant cell type in the vessel wall and are responsible for maintaining vessel wall integrity, elasticity, and contractility (*1*). In healthy vessels, VSMCs maintain a quiescent, contractile phenotype, but in response to injury or aging such as atherosclerosis, VSMCs lose their contractile markers and dedifferentiate into an overproliferative and migratory synthetic phenotype and can transdifferentiate into inflammatory macrophages and calcifying osteoblasts (*2, 3*). Recent lineage-tracing studies have unambiguously demonstrated 30-70% of the cell population in atherosclerotic plaques are of SMC origin and hence, interventions that can modulate VSMC cell fate and phenotype can serve as a powerful strategy to combat atherosclerosis (*4, 5*).

MicroRNAs (miRs) are short, single-stranded non-coding RNAs of 22-24 nucleotide length that demonstrate post-transcriptional gene silencing capabilities (*6*). In particular, miR-145 is the most highly expressed miR in the vasculature and acts as a gatekeeper to VSMCs, regulating and promoting the contractile phenotype (*7, 8*). miR-145 upregulates the contractile genes, myocardin, calponin, and α-SMA, while downregulating synthetic genes by inhibiting Krüppel-like factor-4/5 (KLF-4/5) and ETS domain-containing protein-1 (ELK-1) (*8*). Patients with atherosclerosis have been reported to have reduced levels of circulating miR-145 and in mice, regions of vascular damage express little to undetectable levels of miR-145 (*9–11*). Thus, delivering miR-145 to VSMCs has the potential to be an effective therapy for atherosclerosis.

Key challenges to miR delivery, however, include the lack of nucleic acid stability in serum as well as limited bioavailability to target diseased tissues which can lead to adverse side effects (*12*). These limitations can be overcome with nanoparticle-based delivery systems that protect nucleic acid cargo against serum nucleases and facilitate intracellular uptake for efficient gene therapy (*13*). Most notably, patisiran, an siRNA nanoparticle therapy for hereditary amyloidosis, gained recent FDA approval, bringing much enthusiasm for RNA-based nanotherapies for clinical use (*14*). In addition, several miR-based therapies are in varying phases of pre-clinical and clinical trials for diseases ranging from melanoma to heart failure, highlighting the promise of miR formulations for clinical translation. Nonetheless, many challenges still remain for miR therapy including low drug loading, poor pharmacokinetic properties, and systemic toxicity, limiting its use particularly for diseases such as atherosclerosis that will require a lifetime of repeated drug exposure (*13, 15*). Previously, we described peptide-functionalized micelles that can be designed to target a variety of pathological cell types and features in atherosclerosis including endothelial cells, monocytes, calcification, and microthrombi, and reported their stability and biocompatibility *in vivo* (*16–18*). Given recent data showing VSMCs give rise to the majority of atherogenic cells in plaques upon transdifferentiation, in this study, we aimed to test the suitability of targeting VSMCs as a novel therapeutic strategy against atherosclerosis. We utilize CCR2-targeting micelles to deliver miR-145 to plaques, as pathological VSMCs are marked by the expression of inflammatory cytokines including monocyte chemoattractant protein-1 (MCP-1) as well as its receptor CCR2 (*19*). We show miR-145 micelles efficiently release their nucleic acid cargo upon cellular entry *in vitro* and can rescue the loss of contractile markers in patient-derived SMCs harvested from atherosclerotic arteries. Treatment of ApoE^−/−^ mice in mid-stage atherosclerosis with miR-145 micelles induced downregulation of synthetic markers and upregulation of contractile markers in VSMCs observed through qRT-PCR analyses, and mitigated plaque development *in vivo* confirmed by *en face* analysis and lesion size measurements *via* histology. Notably, miR-145 micelles reduced necrotic core area while increasing collagen content, suggesting miR-145 delivery improved overall plaque stability and decreased likelihood of rupture and risk of acute thrombosis (*20*). Moreover, in early-stage atherosclerotic mice, a single-dose treatment of miR-145 micelles significantly inhibited lesion development by up to 49% in lesion size compared to nontreatment controls. Together, we report the therapeutic potency of delivering miR-145 to intervene during atherosclerosis and a nanocarrier strategy that can be applied at multiple stages of chronic diseases.

## Results

### Synthesis, characterization, cell entry, and intracellular release of miR-145 micelles

To synthesize miR-145 micelles, miR-145 mimics were thiolated on the 5’ end of the sense (functional) strand and covalently conjugated to DSPE-PEG(2000)-maleimide *via* a thioether linkage (**Fig. S1**). miR-145 micelles were constructed with miR-145 amphiphiles, MCP-1 peptide amphiphiles, and DSPE-PEG(2000)-methoxy at a 1:49:50 molar ratio, respectively, to replicate doses used for lentiviral transduction of miR-145 in previous studies and maintain the critical micelle concentration (CMC) needed for micelle self-assembly (**Fig. S2A**) (*7*). miR-145 amphiphile incorporation into micelles did not compromise the size and surface charge of the particle and were similar in particle physicochemical properties as non-therapeutic, MCP-1 micelles (**Fig. S2**). Transmission electron microscopy (TEM) images showed miR-145 micelles were monodispersed, spherical nanoparticles (**Fig. S2B)**, and the hydrodynamic diameter and the surface charge was measured to be 21.7 ± 2.2 nm and 15.1 ± 0.2 mV using dynamic light scattering (DLS) (**Fig. S2F**). We hypothesized that, in addition to PEGylation, this modest positive charge, attributed from the net +6 charge of the MCP-1 peptide, would be favorable for prolonged systemic circulation and intracellular endosomal escape, as opposed to other cationic polymer-based nanoplatforms proposed for nucleic acid delivery which are highly positive (>30 mV) and have reported nonspecific adsorption, particle aggregation, and cytotoxicity (*21*). Control miRs and scrambled, non-targeting peptides were synthesized and fabricated into micelles (miR-67 micelles and NT miR-145 micelles, respectively) and had similar physicochemical properties (**Fig. S2**).

To ensure miR-145 was incorporated into micelles, gel shift assays were conducted, and the mobility of miR-145 micelles was assessed (**Fig. 1A**). Upon electrophoresis, miR-145 micelles remained within the well and did not show migration down the gel (Lane A), indicating miR-145 was fully intact within the micelle at concentrations above the CMC (*22*). In contrast, when miR-145 amphiphiles were loaded at 0.6 µM, a concentration below the 2 µM CMC, as expected, micelles did not assemble and the band containing miR-145 amphiphiles readily migrated down the gel similar to free miR-145 (Lane B and C, respectively). As one of the challenges of effective gene therapy is maintaining the stability of nucleic acids, we also performed gel shift assays in the presence of nucleases to assess the benefit of miR delivery by micelles. miR-145 micelles, miR-145 amphiphiles below the CMC, and free miR-145 were incubated in nuclease-treated FBS for up to 24 hours and migration observed in the gels. As micelles, miR degradation was significantly abated over 24 hours, retaining 89.6 ± 4% of miRs as seen by the strong band intensity within the wells (Lane 3). However, without the protective micellar system, miR-145 amphiphile monomers (below the CMC) and free miR-145 signal migrated down the gel and decreased to 38.2 ± 1% and 29.2 ± 20.8% (Lane 4 and 5) by 24 hours, respectively. Lanes 1, 2, and 6 show untreated miR-145 micelle, untreated miR-145, and FBS only, respectively, which show miRs remain intact without nuclease treatment.

**Fig. 1.**
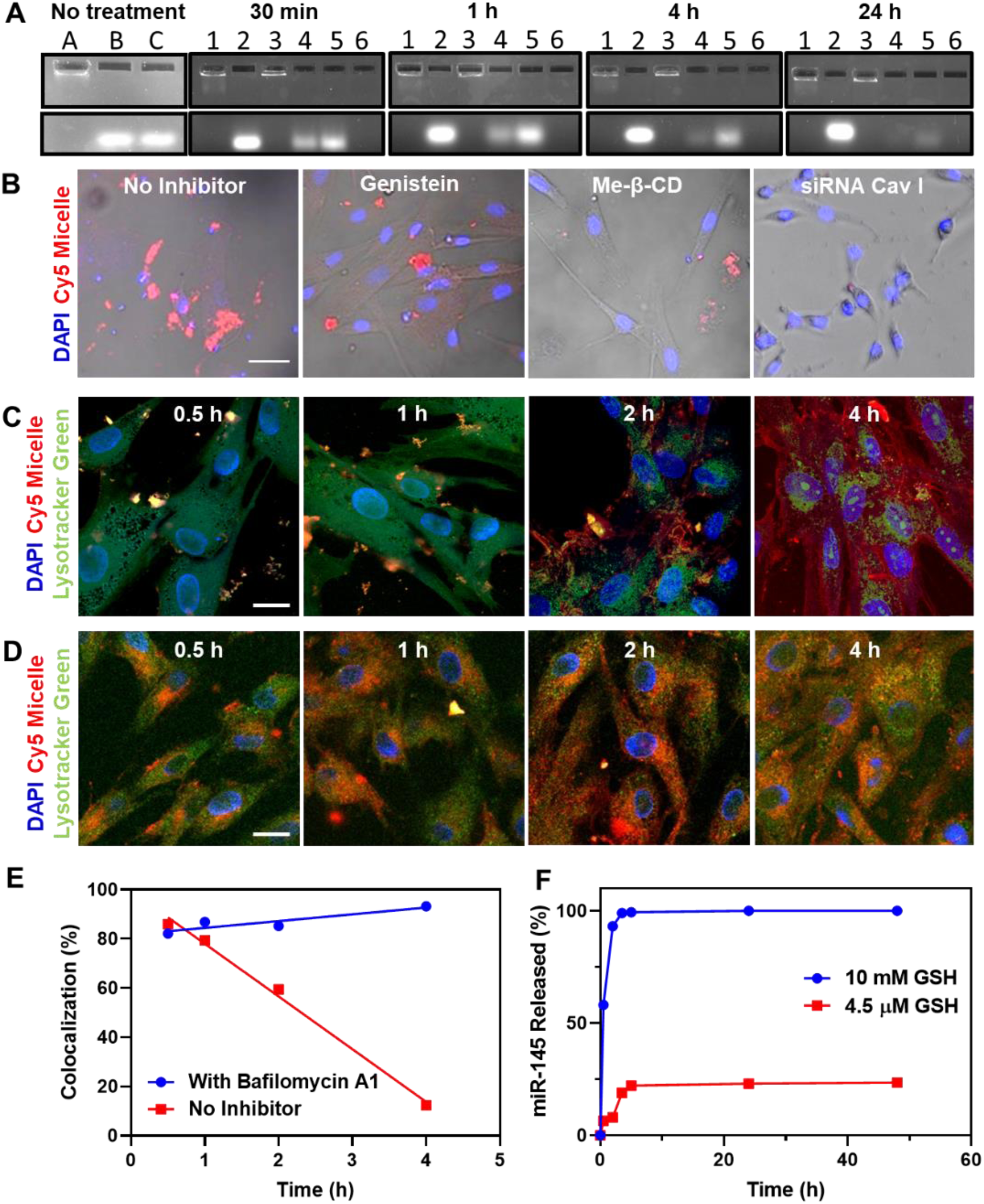
Characterization, cellular internalization, endosomal escape mechanism, and release of miR-145 micelles. (A) Gel shift assays of miR-145 micelles show complete encapsulation of miR-145 within the micelle and reduced degradation under nuclease treatment over 24 hours. Lanes A) untreated miR-145 micelles, B) untreated miR-145 amphiphile below CMC, C) untreated free miR-145. Lanes 1) untreated miR-145 micelle, 2) untreated free miR-145, 3) FBS-treated miR-145 micelle, 4) FBS-treated miR-145 amphiphile below CMC, 5) FBS-treated free miR-145, and 6) FBS. (B) Fluorescence images of hASMCs incubated with Cy5miR-145 micelles (red, Cy5 micelle) with endocytosis inhibitors (energy-dependent (NaN_3_), caveolae-mediated endocytosis (genistein, siRNA caveolin 1), lipid rafts (me-β-CD)). Scale bar = 50 µm. Time-dependent endosomal escape of Cy5miR-145 micelle (red) in hASMCs (C) without and (D) with bafilomycin A1, a vacuolar proton pump inhibitor. Nuclei were stained with DAPI (blue). Scale bar = 20 µm. (E) Percent colocalization of miR and endosomes. (F) Release profiles of miR-145 from micelles show retention of miR-145 at extracellular GSH concentrations (4.5 µM) and release of miR-145 at intracellular GSH concentrations (10 mM).

Next, in order to determine the route of intracellular uptake of micelles, we incubated hASMCs with endocytosis pathway inhibitors including chlorpromazine (clathrin-mediated endocytosis inhibitor), wortmannin (macropinocytosis inhibitor), genistein (caveolin-mediated endocytosis inhibitor), methyl-β-cyclodextrin (lipid raft-mediated endocytosis inhibitor, me-β-CD), and sodium azide (energy-dependent endocytosis inhibitor, NaN_3_, **Fig. 1B, S3**). hASMCs incubated with genistein, me-β-CD, and sodium azide showed a significant decrease in micelle internalization suggesting that miR-145 micelles enter cells *via* energy-dependent, caveolae-mediated endocytosis lipid rafts. This was further confirmed when miR-145 micelle uptake decreased after hASMCs were transfected with siRNA caveolin I (**Fig. 1B**) and micelles showed colocalization with cholera toxin subunit B (CT-B) Alexa Fluor488, a protein known for internalization *via* caveolin-mediated endocytosis **(Fig. S4)**.

Next, to test endosomal escape of miR-145 micelles after cell internalization, hASMCs were incubated with Cy5-labeled, miR-145 micelles (Cy5 micelle) for up to 4 hours and stained with Lysotracker Green. Internalization was observed at 30 minutes, confirmed by the co-localization of micelles (red) and endosome (green, **Fig. 1C**), and by 4 hours, a linear decrease in co-localization was seen, indicating miR-145 micelles successfully escaped the endo-lysosomal compartment and premature degradation. Similar to other cationic nanoparticles, the successful endosomal escape of miR-145 micelles may be attributed to the proton sponge effect (*23*). This was verified through the treatment of hASMCs with miR-145 micelles in the presence of a vacuolar proton pump inhibitor, bafilomycin A1, which prevents the acidification of lysosomes. As shown in **Figure 1D**, incubation with bafilomycin A1 led to the retention of miR-145 micelles within the endosome/lysosome, determined by the high co-localization (82%-94%) between endosome/lysosome (green) and micelles (red) over time (**Fig. 1E**). To further confirm this mechanism of endosomal escape, we analyzed the downstream effect of miR-145 micelles in hASMCs upon bafilomycin A1 incubation. **Figure S5** confirmed that the synthetic (KLF-4, ELK-1) and contractile (α-SMA) markers remained similar between PBS and miR-145 micelle treatments in the presence of bafilomycin A1, but without the inhibitor, miR-145 micelles decreased synthetic markers and increased contractile marker (**Fig. S5**).

Once in the cytoplasm, our initial data (**Fig. S5)** suggested that the thioether linkage between miR-145 and PEG is cleaved and miR-145 is successfully released from nanoparticles to induce contractile SMC gene expression. As the cytosolic level of glutathione (GSH) is significantly higher than extracellular and endosomal GSH levels, GSH can act as a natural reducing agent for the release of cargo from nanoparticles (*24*). Thus, covalent linkage would prevent the premature release of miR-145 in the low extracellular GSH concentration (4.5 µM) until it can be reduced by the elevated GSH concentration (10 mM) upon reaching the cytosol (**Fig. 1F**) (*24, 25*). While only 19% of miR-145 was released from miR-145 micelles in environments with 4.5 µM GSH, at cytosolic levels of GSH (10 mM), 100% of miR-145 was released by 4 hours confirming cleavage of miR-145 under cytosolic GSH concentrations and the thioether linkage to hinder premature miR release prior to cell internalization.

### Modulation of atherosclerotic genes *in vitro* and in VSMCs derived from patients with cardiovascular disease

After validation of successful miR-145 micelle uptake and transfection, we investigated the efficacy of miR-145 micelles to modulate VSMC phenotype and its potential as a therapy for atherosclerosis. As miR-145 prevents plaque development through the elevation and inhibition of genes responsible for the contractile and synthetic SMC phenotypes, respectively, we incubated hASMCs with miR-145 micelles and evaluated SMC markers through qRT-PCR (*8*). As shown in **Figure 2A**, miR-145 micelles downregulated athero-prone KLF-4/5 and ELK-1 expression (KLF-4: 44.3 ± 5.3%, KLF-5: 56.5 ± 6.2%, ELK-1: 30.4 ± 1.8%, **Fig. 2A)** and simultaneously upregulated the athero-protective genes, myocardin (162.1 ± 10.7%), α-SMA (166.7 ± 12.5%), and calponin (142.5 ± 18.2%) relative to PBS treatment groups (**Fig. 2B**). Consistent with other reports on miR-145, miR-145 micelles also downregulated CCR2 expression to 58.2 ± 4.3% relative to PBS treatment groups, which is normally overexpressed in synthetic VSMCs (**Fig. 2C**) (*26*). In contrast, non-therapeutic control miR-67 micelles did not induce any effects on the targeted genes.

**Fig. 2.**
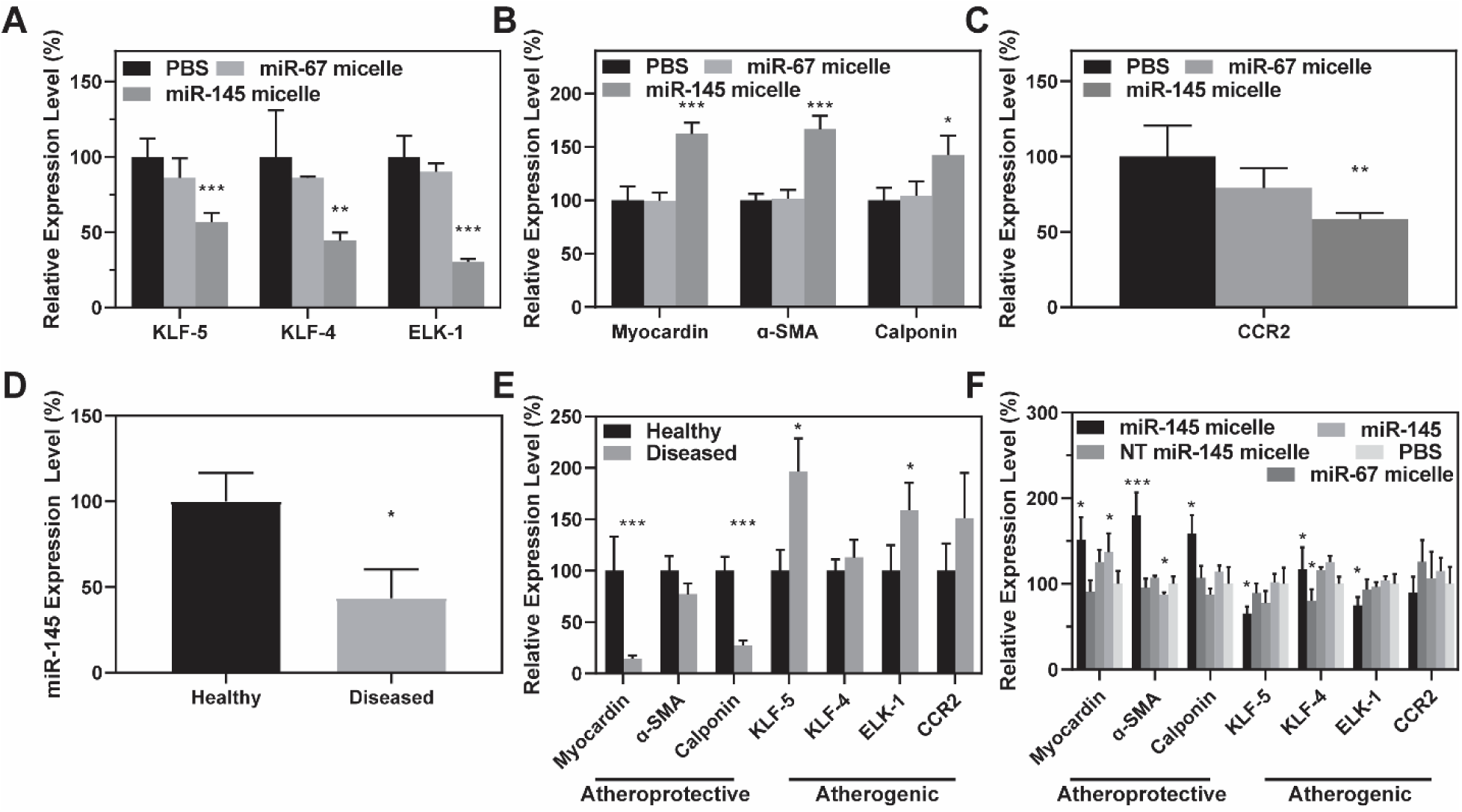
miR-145 micelles silence synthetic markers and rescue contractile markers in VSMCs. mRNA expression of hASMCs transfected with miR-145 micelles show downregulation of (A) synthetic SMC markers, upregulation of (B) contractile SMC markers, and decreases in (C) CCR2 (N = 3; *p < 0.05, **p < 0.01, ***p < 0.001 relative to PBS). Patient-derived SMCs isolated from arteries with atherosclerotic plaques have (D) lower miR-145 expression compared to SMCs from healthy tissue (N = 3; *p < 0.05) and (E) lower endogenous expression of contractile markers and higher expression of synthetic markers (N = 4; *p < 0.05, ***p < 0.001 compared to healthy samples). (F) miR-145 micelle treatment on VSMCs derived from diseased patient tissue rescue contractile markers and downregulate synthetic markers (N = 4; *p < 0.05, ***p < 0.001 relative to PBS).

To further assess the therapeutic potential of miR-145 micelles, SMCs were isolated from discarded tissue from both atherosclerotic and healthy arteries from patients undergoing vascular surgery. First, we found that endogenous miR-145 was 56.4 ± 16.8% lower in SMCs derived from diseased arteries compared to miR-145 in SMCs from normal arteries (**Fig. 2D**). In addition, contractile markers, myocardin (14.2 ± 3.2%) and calponin (27.2 ± 5.0%), were downregulated, while synthetic markers, KLF-5 (196.9 ± 31.8%) and ELK-1 (158.8 ± 26.6%), were overexpressed in VSMCs from diseased tissue compared to healthy tissue (**Fig. 2E**). No statistical differences were observed for α-SMA, KLF-4, and CCR2 expression levels, possibly due to location differences of VSMC origin (e.g. tibial artery vs. aorta), extent of disease progression in these tissues (e.g. moderate vs. severe atherosclerosis), and small sample size.

Patient-derived VSMCs from diseased tissue were then treated with miR-145 micelles and phenotypic markers were analyzed *via* qRT-PCR (**Fig. 2F**). Consistent with **Figures 2B** and **C**, when incubated with miR-145 micelles, the expression of contractile markers myocardin (151.8 ± 26.0%), α-SMA (180.3 ± 26.5%), and calponin (158.8 ± 21.6%) increased, while synthetic markers KLF-5 (65.3 ± 8.2%) and ELK-1 (74.9 ± 9.8%) were significantly decreased relative to PBS treatment groups. NT miR-145 micelles and free miR-145 also showed an increase in myocardin expression, but no statistical differences in α-SMA and calponin, as well as KLF-5, ELK-1, and CCR2 were found compared to PBS controls. These results demonstrate that micelle nanoparticles enhance miR-145 uptake by VSMCs and can rescue the contractile phenotype of synthetic SMCs derived from diseased arteries.

### *In vivo* pharmacokinetics, biodistribution, and targeting of miR-145 micelles

To assess the stability of miR-145 micelles *in vivo*, the pharmacokinetic properties of miR-145 micelles co-assembled with Alexa Fluor 647 (AF647)-labeled miR-145 amphiphiles, Cy7 amphiphiles, and MCP-1 amphiphiles were investigated in wild-type C57BL/6J mice. After 24 hours post-intravenous administration, the circulation half-lives were fitted into a two-compartment model. While the overall biodistribution profiles of miR-145 micelles were in accordance with other micellar systems (**Fig. S6**) importantly, Cy7 (micelle) and AF647 (miR) blood circulation half-lives (t_1/2β_) were found to be similar (10.3 ± 3.1 hours and 14.3 ± 2.6 hours (**Fig. 3A**), respectively), and not statistically significant (p = 0.1016) (*22*). Additionally, the ratio of Cy7 amphiphiles and miR-145 remained consistent up to 24 hours post-injection, suggesting that miR-145 remained assembled within the micelle during systemic circulation. In contrast, free miRs are well known to be rapidly cleared by the reticuloendothelial system and the kidneys, with reported half-life of 30 minutes, which highlights the importance of the protective, micelle carrier to provide prolonged availability of nucleic acids *in vivo* (*27*). Hematoxylin and eosin (H&E) staining of all organ sections showed no obvious toxicity, and the morphology of organs was similar among PBS-, miR-145 micelle-, and NT miR-145 micelle-treated mice (**Fig. S7A**). Moreover, blood urea nitrogen and creatinine levels were similar among all groups and within the range of healthy C57BL/6J mice, confirming kidney health and safety of these particles *in vivo* (**Fig. S7B**).

**Fig. 3.**
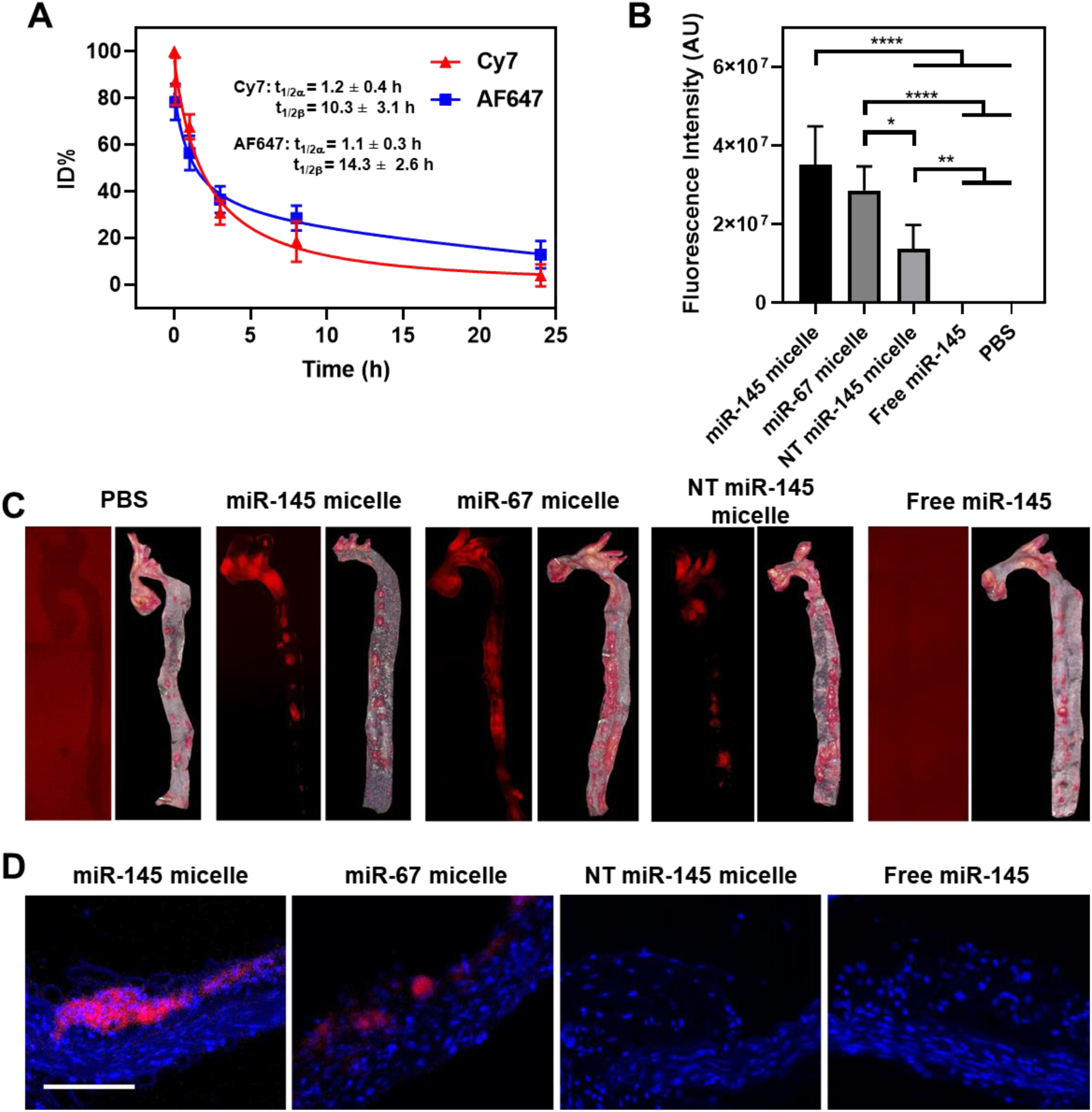
Pharmacokinetics and *in vivo* targeting of miR-145 micelles. (A) Half-life of Cy7 (micelle) and AF647 (miR-145) in blood using a two-compartment model were similar, indicating miRs remained assembled in micelles (N = 4). (B) Fluorescence quantification in aortas of atherosclerotic mice show miR-145 micelles accumulate to the greatest degree compared to miR-67 micelles and NT miR-145 micelles (N = 6). (C) Whole aorta fluorescence imaging shows micelles target plaques identified by ORO. (D) Histological analyses show miR-145 micelles (red) accumulate in plaques. Scale bar = 100 µm.

Targeting efficiency of miR-145 micelles was then tested in murine models of atherosclerosis. Specifically, ApoE^−/−^ mice were fed a high-fat diet for 14 weeks to develop mid-stage atherosclerosis before treatment with five injections, every three days of PBS, miR-145 micelle, miR-67 micelle, NT miR-145 micelle, or free miR-145 (*28*). Twenty-four hours after the last injection and 17 total weeks of high-fat diet, aortas were harvested and imaged before histological assessment (**Figs. 3B, 3C, 3D**). *Ex vivo* fluorescence imaging of whole aortas from mice treated with miR-145, miR-67, or NT miR-145 micelles showed particles localized to regions of plaques identified *via* an oil red o (ORO) lipid stain (**Figs. 3C**). This result is further corroborated by histological analyses of artery cross-sections that showed the Cy7 particle signal confined to atherosclerotic plaques (**Fig. 3D**). miR-145 micelles (3.5 × 10^7^ ± 9.9 × 10^6^ AU) and miR-67 micelles (1.4 × 10^7^ ± 6.0 × 10^6^ AU) demonstrated the highest signal upon quantification, followed by NT miR-145 micelles (2.9 × 10^7^ ± 6.1 × 10^6^ AU), indicating that MCP-1 peptides facilitated enhanced plaque targeting (**Fig. 3B**). Some nonspecific diffusion of NT miR-145 micelles into the plaques can be attributed to endothelium disruption that is commonly observed in atherosclerotic plaques (*29*). However, no fluorescence signal was detected in the aorta of mice treated with free miR-145 and was similar to PBS controls (**Figs. 3B, 3C**, and **3D**). Importantly, there was no loss in body weight in all mice, indicating the nanoparticle dosages were well-tolerated during the three weeks of treatment (**Fig. S7C, S7D**).

### Atherosclerosis inhibition by miR-145 micelles in advanced atherosclerosis *in vivo*

Atherosclerosis is often an asymptomatic disease with undetected vulnerable plaques that can rupture precipitously, causing acute cardiac events. Thus, we tested the therapeutic potential of miR-145 micelles to reduce and stabilize atherosclerosis in ApoE^−/−^ mice that were fed a high fat diet for 14 weeks that develop mid-stage disease. At week 14, mice in treatment groups were injected once every three days for a total of five injections while simultaneously fed a high fat diet (17 weeks total high fat diet). When total plaque area in the aortas were evaluated using *en face* ORO stain (**Fig. 4A** and **4B)**, mice treated with miR-145 micelles showed the least amount of plaques (20.0 ± 4.5% of plaques to total aorta surface area) compared to all other control groups (miR-67 micelle: 29 ± 5.9%; p = 0.0014 vs. NT miR-145 micelle: 35.6 ± 10.3%; free miR-145: 25.4 ± 4.7%; p = 0.0004 vs. PBS: 40.0 ± 6.4%). As shown in **Figures 4C** and **4D**, immunohistochemistry and qRT-PCR analyses of aortic root plaque lesions demonstrated that mice treated with miR-67 micelles, NT miR-145 micelles, or free miR-145 showed similar expression levels of contractile markers (77-123%), synthetic markers (83-122%), and miR-145 expression (78-100%) relative to PBS. In contrast, miR-145 micelle treatment showed a significant increase in miR-145 expression in plaques (140.3 ± 21.6% vs. 192.7 ± 60.8% for miR-145 micelles vs. baseline) and upregulation of contractile markers that were similar to baseline groups sacrificed at week 14 (myocardin: 145.4 ± 20.4% vs. 151.6 ± 17.2%; α-SMA: 200.4 ± 43.9% vs. 221.4 ± 24.4%; calponin: 167.8 ± 24.7% vs. 177.2 ± 18.3% for miR-145 micelles vs. baseline all relative to PBS), as well as downregulation of synthetic markers (KLF-5: 63.1 ± 11.8% vs. 68.3 ± 11.1%; KLF-4: 83.6 ± 8.1% vs. 92 ± 10.8%; ELK-1: 69.6 ± 9.2% vs. 71.6 ± 31.2% for miR-145 micelles vs. baseline all relative to PBS; **Figs. 4C** and **D**). Immunohistochemistry confirmed the increase in calponin and α-SMA expression at the lesion site compared to miR-67 micelles, NT miR-145 micelles, miR-145, and PBS which showed minimal expression, further supporting the therapeutic benefit of miR-145 micelles in mitigating plaque growth or atherosclerosis progression (**Fig. S8**).

**Fig. 4.**
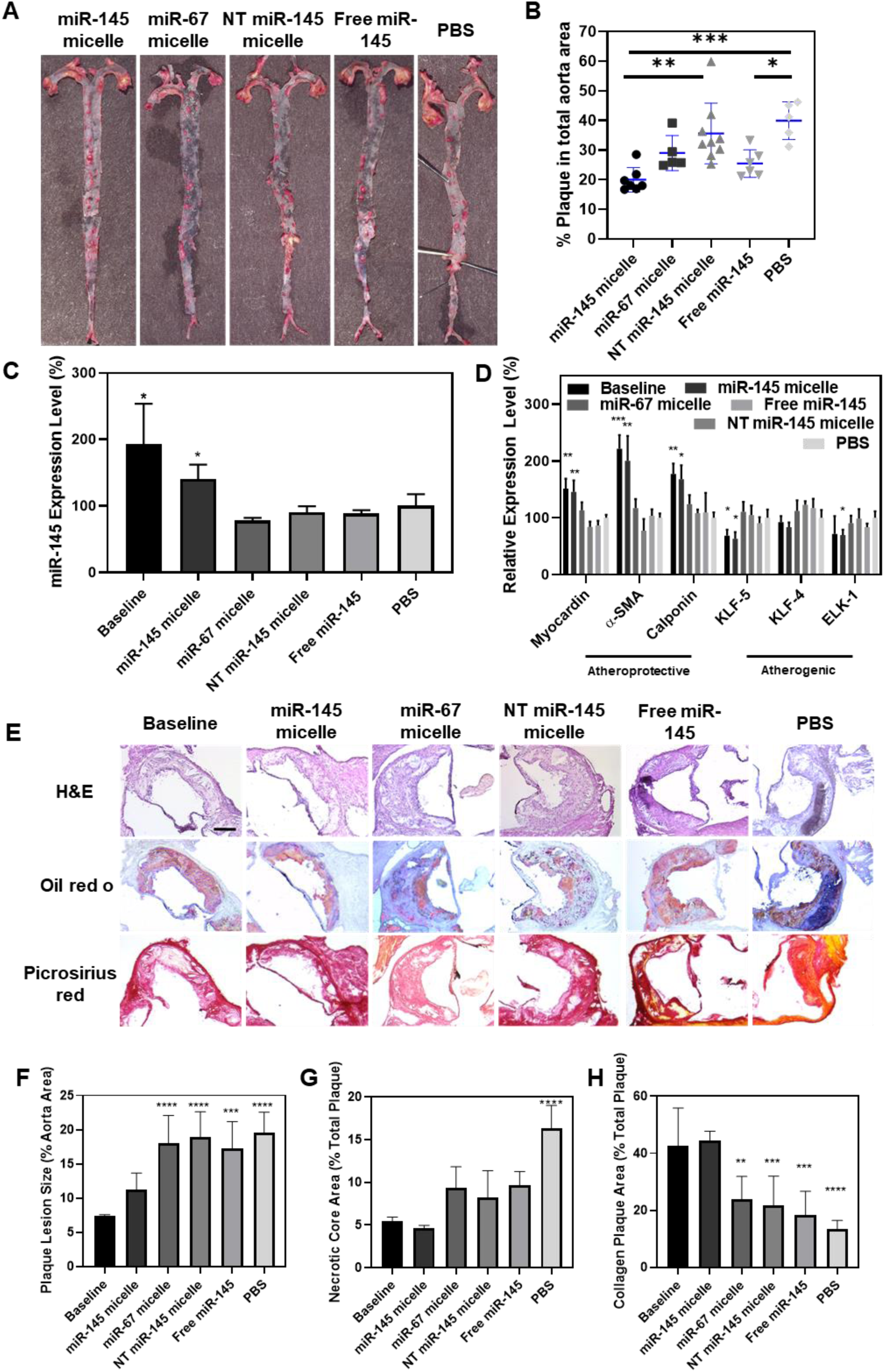
miR-145 micelles rescue the SMC contractile phenotype and reduce plaque progression *in vivo*. (A) ORO *en face* staining show fewer lesions in the aortic arch and descending aorta in miR-145 micelle-treated mice. (B) Quantification of ORO staining shows lower total plaque area upon miR-145 micelle treatment (N > 5). (C) miR-145 expression and (D) mRNA expression of contractile and synthetic phenotype markers in VSMCs from mid-stage atherosclerotic mice (N = 6; *p < 0.05, **p < 0.01, ***p < 0.001 relative to PBS). miR-145 micelles increase miR-145 expression and contractile markers, while decreasing synthetic markers. (E) Representative histological images of the aortic root stained with H&E, ORO, and picrosirius red (collagen) staining. Scale bar = 200 µm. Quantification of (F) plaque lesion size, (G) necrotic core area, and (H) collagen (N > 11 for treatment groups, N = 6 for baseline; *p < 0.05, **p < 0.01, ***p < 0.001 as compared to baseline).

Through histological analyses, the increase in miR-145 and SMC contractile markers found in miR-145 micelle-treated mice correlated with a decrease in atherosclerotic plaque lesion size and necrotic core area by 43% and 72%, respectively (vs. PBS-treated controls, p < 0.0001, **Figs. 4E, F** and **G**). Decreases in lesion size may be attributed to fewer synthetic and macrophage-like SMCs that form the lipid necrotic core (*30*). To confirm this, we analyzed the aorta of treated mice for CD107b+ macrophage markers and found that miR-145 micelle treatment decreased CD107b+ signal compared to other treatment groups (**Fig. S8**). Furthermore, CCR2 expression associated with inflammation and synthetic SMCs in the aortic root was similar between the miR-145 micelle treatment group and baseline (**Fig. S9A**). Together, miR-145 micelle treatment contributed to the reduction of necrotic cores and plaque size to mitigate atherosclerosis.

Notably, while synthetic SMCs typically release matrix metalloproteinases (MMPs) that degrade collagen, contractile SMCs within plaques are capable of synthesizing structurally enhancing collagen isoforms that provide plaque stability through the thickening of the fibrous cap (*31*). Hence, miR-145 micelles may increase collagen abundance and stabilize plaques due to their ability to rescue contractile SMCs. To assess this, we analyzed plaques of treated mice and found miR-145 micelle treatment had the most abundant intraplaque collagen (44.6 ± 3.1% collagen area of total plaque area vs. PBS: 13.5 ± 2.9%, p < 0.0001, **Fig. 4H**), while mice in PBS, miR-67 micelle, NT miR-145, and free miR-145 treatment groups developed larger plaques, exhibited increased necrotic core area, and had lower collagen content in the fibrous cap which are characteristic to plaques that are rupture-prone (*20*). No statistical differences were found among miR-67 micelles, NT miR-145 micelles, free miR-145, and PBS treatment groups regarding plaque lesion size, necrotic core area, and collagen content. These initial results indicate miR-145 micelle treatment is a viable and safe therapy for prolonged treatment of atherosclerosis and future studies will assess long-term safety of miR-145 micelles.

### Atherosclerosis prevention by miR-145 micelles *in vivo*

Lineage tracing studies have shown that VSMCs can adopt phenotypes related to early markers of atherosclerosis including foam cells, macrophages, and synthetic SMCs (*30*). Hence, we hypothesized that early dosing of VSMCs with miR-145 micelles could mitigate early plaque formation by preventing the formation of fatty streaks and lesions attributed to foam cells, decreasing inflammation contributed by macrophages, and inhibiting overproliferation of synthetic SMCs. To test this hypothesis, we first examined the cholesterol efflux capabilities of cholesterol-loaded and activated hASMCs and found that miR-145 micelles induced 80% cholesterol efflux (**Fig. S9B**). In addition, to test this *in vivo*, ApoE^−/−^ mice were fed a high-fat diet for four weeks to develop early-stage atherosclerosis. After two weeks of high-fat diet representing a “pre-disease state,” mice were treated with a one-time dose of miR-145 micelles, miR-67 micelles, NT miR-145 micelles, free miR-145, or PBS. By the end of four weeks, VSMCs were isolated and subjected to qRT-PCR and histological analyses. As shown in **Figure 5A**, strikingly, miR-145 micelles increased expression of miR-145 to 178.5 ± 22.9%, whereas no change in miR-145 expression was observed for mice treated with miR-67 micelles, NT miR-145 micelles, and free miR-145 relative to PBS (p < 0.01). Notably, the increased expression of miR-145 provided by miR-145 micelles induced contractile SMC gene expression (179.4 ± 25.5%, p < 0.05; 141.9 ± 7.9%, p < 0.01; and 180.1 ± 11.4%, p < 0.001; for myocardin, α-SMA, and calponin, respectively, vs. PBS, **Fig. 5B**). Likewise, miR-145 micelle treatment reduced synthetic SMC marker expression including KLF-5, KLF-4, and ELK-1 to 58.0 ± 7.0% (p < 0.01), 60.2 ± 8.2% (p < 0.01), and 53.2 ± 3.6% (p < 0.001), respectively. In contrast, mice treated with miR-67 micelles, NT miR-145 micelles, and free miR-145 exhibited no statistical differences in expression of SMC contractile or synthetic phenotype markers relative to PBS-treated controls. These results strongly suggest that early dosing of VSMCs with miR-145 micelles can prevent SMCs from adopting alternative phenotypes that cause propagation of atherosclerotic plaques.

**Fig. 5.**
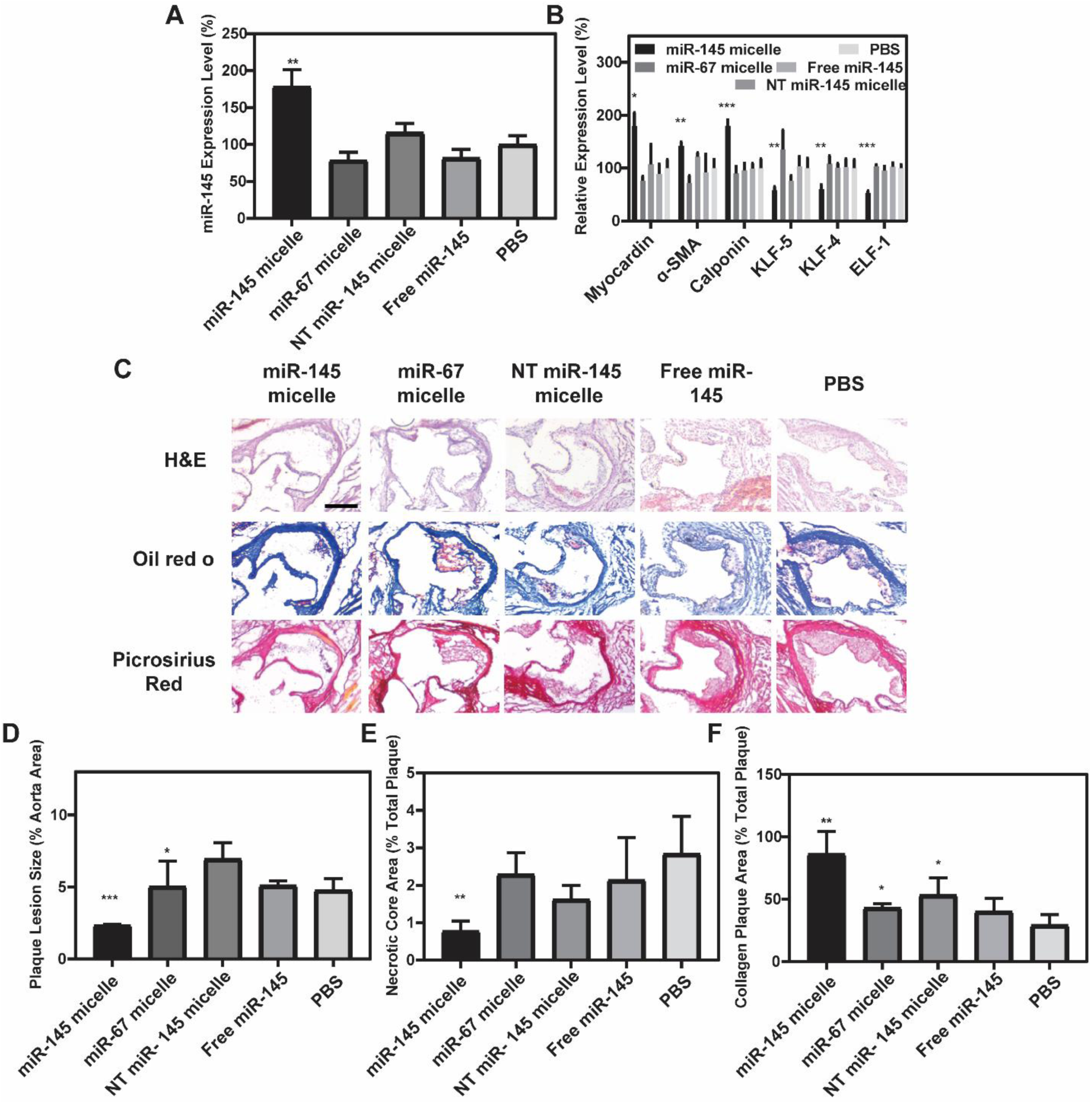
miR-145 micelles promote contractile SMC phenotype and prevent atherogenesis in early-stage disease. (A) miR-145 expression and (B) mRNA expression of myocardin, α-SMA, calponin, KLF-5, KLF-4, ELK-1 analyzed in VSMCs from early-stage atherosclerotic mice (N = 6; *p < 0.05, **p < 0.01, ***p < 0.001 relative to PBS). (C) Representative histological images of the aortic root stained with H&E, ORO, and picrosirius red. Scale bar = 200 µm. Quantification of (D) plaque lesion size, (E) necrotic core area, and (F) collagen containing plaque area demonstrate significant reduction in lesion size and necrotic core area with increases in collagen after miR-145 micelle treatment (N = 6; *p < 0.05, **p < 0.01, ***p < 0.001 compared to PBS).

Upon histological analysis, mice treated with miR-145 micelles showed a reduction in lesion size and necrotic core area by 49 ± 0.1% (p < 0.001) and 73 ± 0.3% (p < 0.01), respectively, compared to the PBS-treated control group (**Figs. 5C, 5D**, and **5E**). miR-145 micelle treatment significantly enhanced collagen content within the fibrous cap of atherosclerotic plaques by 292% (p < 0.01). NT miR-145 micelles and miR-67 micelles exhibited some increases in collagen content within these lesions as well, which is likely attributed to diffusion of NT miR-145 micelles to plaques as seen in **Figure 3D** (**Fig. 5F**). Therefore, these results suggest that miR-145 micelles increase α-SMA-positive, contractile SMCs that promote plaque stability. Furthermore, miR-145 micelles caused a significant decrease in CCR2 expression and serum inflammatory cytokines (p < 0.01 vs. PBS for TNF-α, **Fig. S9C, S10**) and a decrease in pro-inflammatory cytokine IL-6 (p < 0.05 vs. PBS, **Fig. S10**) to levels similar to that of C57BL/6J mice on a standard diet as shown in previous studies (*32*). In addition to decreasing inflammation and disease progression, the minimal inflammatory response to miR-145 micelles indicates strong tolerability of this nanotherapy, favoring extended nanoparticle treatment for chronic disease.

## Discussion

Atherosclerosis is a complex inflammatory disease that can take decades to progress. Treatments for atherosclerosis must consider strategies that can intervene at early as well as late-stage disease, especially for high-risk populations and those with a family history of cardiovascular disease. Despite the involvement of VSMCs throughout atherogenesis, and the prevalence of macrophage-like and osteoblast-like cells of VSMC origin in atherosclerotic plaques, few studies have demonstrated the potential of modulating VSMC phenotype as a therapeutic strategy for atherosclerosis. miR-145 has been identified as a key regulator in promoting SMC contractile phenotype by inhibiting KLF-4 and promoting myocardin (*7*). Thus, delivering miR-145 has the potential to significantly limit plaque development and growth in atherosclerosis.

Our studies demonstrate miR-145 delivery by micelles slows atherosclerosis progression. We first show that miR-145 micelles are efficiently internalized into VSMCs and escape the endo-lysosomal compartment to release the miR-145 payload in the cytosol, promoting the contractile phenotype *in vitro* (**Fig. 1**). Of significance, we show miR-145 micelles have clinical relevance by rescuing contractile SMC markers, myocardin, α-SMA, and calponin, in patient-derived, diseased VSMCs that had a loss of miR-145 expression (**Figs. 2D, 2E, 2F**). As previous studies report lower circulating levels of miR-145 in atherosclerotic patients and within their plaques, our early results using patient-derived VSMCs show potential for clinical translation of miR-145 (*10, 33*). CCR2-targeting peptides were included into miR-145 micelles as CCR2 is upregulated in synthetic VSMCs associated with atherosclerosis as well as inflammatory atherogenic cells found in plaques (*26*). However, as CCR2 is also expressed on endothelial cells and monocytes involved in atherosclerosis (*18*), future studies will need to elucidate the potential therapeutic contribution upon uptake of miR-145 micelles in these cell types.

Importantly, miR-145 delivery by micelles enhanced efficacy in reducing plaques compared to free miR-145 and non-targeting micelles upon *en face* and histological analyses (**Figs. 4** and **5**). The targeting moiety to CCR2, near-neutral surface charge and stability that significantly prolongs blood circulation *in vivo*, as well as the efficient cell internalization and endosomal escape of miR-145 micelles contribute to these results (**Fig. 3**). One important observation that will need to be fully addressed in future studies is the dramatic effects of a single dose of miR-145 micelles in altering miR-145 expression and reducing lesion size in early, developing atherosclerotic plaques (**Fig. 5**). Recent studies with VSMC lineage tracing have shown that the clonal expansion of a few VSMCs in the arterial wall are responsible for most VSMC-derived cells in plaques (*34*). Hence, early intervention of VSMC proliferation and transdifferentiation could induce an amplified effect in inhibiting atherosclerosis plaque development (*8*). In addition, these results may be explained by the enhanced permeability and retention effect that is more prominent in early, developing atherosclerosis compared to late-stage plaques (*35*). Interestingly, inclisiran, an investigational siRNA-based therapy in clinical trials as a PCSK9 inhibitor for atherosclerosis, has also shown a single dose to have long-term benefits in humans for up to a year (*36*). Overall, our results demonstrate miR-145 therapy as a viable and safe strategy for treating atherosclerosis, and the benefit of infrequent dosing may be used to supplement patients who are intolerant of current first-line, atherosclerosis therapies.

## Materials and Methods

### General Experimental

All starting materials were purchased from Sigma-Aldrich (St. Louis, MO) and Fisher Scientific (Waltham, MA), unless otherwise noted.

WEHI 271.1 (ATCC# CRL-1679) murine monocytes and hASMCs (ATCC# PCS-100-012, Manassas, VA) were purchased from ATCC and subjected to mycoplasma testing. WEHI 274.1 was cultured in DMEM medium (Thermo Fisher Scientific, Waltham, MA), supplemented with 10% FBS, 0.05 mM 2-mercaptoethanol, and 1% penicillin-streptomycin. hASMCs were cultured in Medium 231 (Thermo Fisher Scientific, Waltham, MA), supplemented with smooth muscle growth supplement (SMGS, Thermo Fisher Scientific, Waltham, MA) and 1% penicillin-streptomycin (Thermo Fisher Scientific, Waltham, MA).

### Synthesis of peptide amphiphiles

MCP-1 and scrambled peptides were conjugated onto DSPE-PEG(2000)-maleimide using a previously reported protocol developed by our lab. Briefly, 0.25 mmol MCP-1 peptide [YNFTNRKISVQRLASYRRITSSK] or scrambled peptide [YNSLVFRIRNSTQRKYRASIST] were synthesized on an automated peptide synthesizer (PS3, Protein Technologies, Tucson, AZ) on Wang resin via standard Fmoc-mediated, solid phase peptide synthesis methods. To cleave the peptide from the resin, a solution of 94:2:5:2:5:1 vol% trifluoroacetic acid:1,2-ethanedithiol:water:triisopropylsilane was added to the beads and reacted for 4 hours. Peptides were dried with nitrogen, precipitated, and washed twice with ice-cold diethyl ether. The crude peptides were dissolved and lyophilized.

MCP-1 and scrambled peptides were dissolved in milliQ water and purified on a reverse-phase high performance liquid chromatography system (HPLC, Prominence, Shimadzu, Columbia, MD) using a Luna C8 column (250 × 10 mm ID, 5 µm, Phenomenex, Torrance, CA) and a gradient mobile phase consisting of (A) 0.1% (v/v) formic acid in water and (B) 0.1% (v/v) formic acid in acetonitrile. The initial mobile phase consisted of 100% solvent A at a flow rate of 10 mL/min. Solvent A was decreased slowly to 0% over 27 minutes and held at this composition for 3 minutes. The gradient was returned to 100% solvent A over 3 minutes and held at this composition for 1 minute to give a total run-time of 34 minutes. The column temperature was kept at 55°C. The m/z was characterized by MALDI-TOF/TOF (Bruker, Billerica, MA, USA). The expected m/z peak is [M+H]+ = 2890 (**Fig. S1**).

The purified peptides were conjugated to DSPE-PEG(2000)-maleimide in pH 7.4 buffered water for 24 hours. The resulting products were purified using a C4 Jupiter column (250 × 30 mm ID, 5 µm, Phenomenex, Torrance, CA) as described above. The expected m/z peak for the peptide amphiphiles are [M+H]+ = 5830 (**Fig. S1**). Fluorescently-labeled (Cy5, Cy7) amphiphiles were synthesized similarly using DSPE-PEG(2000)-amine through an NHS ester reaction.

### Synthesis of DSPE-PEG(2000)-miR-145 mimics

miR-145 containing the sense sequence, 5’-GUCCAGUUUUCCCAGGAAUCCCU-3’, and control miR (miR-67), containing the sense sequence, 5’-UCACAACCUCCUAGAAAGAGUAGA-3’, were custom ordered from IDT (Coralville, IA), and modified with a thiol group on the 5’ end of the sense (functional) strand for covalent conjugation to the micelle lipid tail. miR-145-SH (MW = 14490 g/mol, 117.5 nmol, 1.70 mg) was added to DEPC-treated water to make 0.1 mM miR-145 solution. TCEP was added to the miR-145 solution and stirred in the dark at room temperature for 4 hours at 1600 rpm. Thiolated miR-145 was conjugated to DSPE-PEG(2000)-malemide (Avanti Polar Lipids, Alabaster, AL) *via* a thioether bond by adding a 10% molar excess of lipid to reduced thiolated miR in DEPC-treated water. The resulting products were characterized using MALDI. The expected m/z peaks for DSPE-PEG(2000)-miR-145 and DSPE-PEG(2000)-miR-67 were [M+H]+ = 17047 and 17717, respectively (**Fig. S1**).

### Construction and characterization of miR-containing micelles

miR-145 micelles were self-assembled by first dissolving peptide amphiphiles and DSPE-PEG(2000)-methoxy (49:50 mol ratio) in methanol. Methanol was completely evaporated under nitrogen and further vacuum dried overnight. The resulting film was hydrated with 100 µL of the solution of DSPE-PEG2000-miR-145 (1 mol%) in nuclease-free water or PBS, and incubated at 60°C for 30 min. After incubation, the micelle solution was cooled to room temperature prior to use. Micelles containing miR-67 or scrambled MCP-1 sequence were synthesized in a similar manner.

The particle size and zeta potential of micelles at 100 µM were determined by a Mobius Zetasizer (Wyatt, Santa Barbara, CA), and measurements were carried out in 25°C in three replicates. TEM was used to observe the morphology of miR-145 micelles. Briefly, 100 µM micelle solutions in water were placed on carbon grids (Ted Pella, Redding, CA) and stained with 2 wt% uranyl acetate. Dried samples were imaged on JEM 2100-F (JEOL Ltd., Tokyo, Japan).

### miR-145 micelle protection against nucleases

miR-145 micelles, miR-145 amphiphiles (below CMC), or free miR-145 containing 10 µg miR were incubated with 20 µL nuclease-treated FBS for 0.5, 1, 4, and 24 hours at 37°C. The integrity of miR-145 was observed by gel electrophoresis on a 2% (w/v) agarose gel at 56 V for 1 hour. Lanes were loaded with miR-145 micelles, miR-145 amphiphiles (below the CMC), or free miR-145 at 10 µg/mL. The gel was imaged on a ChemiDoc XRS+ (Bio-Rad Laboratories, Hercules, CA). The intensity of each band was quantified via ImageJ and normalized against untreated miR-145 micelles or free miR-145.

### *In vitro* micelle internalization mechanism

hASMCs were seeded onto glass coverslips in a 6-well plate for 24 hours at 37°C and 5% CO_2_. Cells were incubated with endocytosis and cell uptake inhibitors, including chlorpromazine (10 μg/mL, clathrin-mediated endocytosis inhibitor), genistein (200 μg/mL, caveolae-mediated endocytosis inhibitor, Me-β-CD (50 μM, lipid-raft-mediated endocytosis inhibitor), wortmannin (50 nM, macropinocytosis inhibitor), and NaN_3_ (10 mM, energy-dependent endocytosis inhibitor), for 30 minutes before fresh media replacement, and cells were washed with PBS and incubated with Cy5miR-145 micelles for 4 hours at 37°C. Cells were transfected with siRNA caveolin 1 (Thermofisher, Waltham, MA) using Lipofectamine 2000 (Invitrogen, Carlsbad, CA) following manufacturer’s protocol. Briefly, Lipofectamine reagent was added to Opti-MEM Medium in which siRNA caveolin 1 was diluted. This mixture was incubated with cells for 72 hours before Cy5miR-145 micelle uptake was measured. Cells were stained with DAPI (10 μg/mL) according to manufacturer’s instructions and imaged on a Confocal Laser Scanning Microscope 780 (Zeiss, Oberkochen, Germany) at an excitation wavelength of 405 nm and 650 nm to visualize the nucleus (blue) and the micelle internalization (red), respectively.

### *In vitro* release profiles

To evaluate the release of miR-145, miR-145 micelles consisting of 1 µg AF647-labeled miR-145 were incubated in PBS at 37°C while stirring and supplemented with 4.5 µM GSH or 10 mM GSH to stimulate the extracellular and intracellular reducing environments, respectively. At 0, 0.5, 2, 3.5, 5, 24, and 48 hours, 5 µL of samples were injected in the HPLC (Prominence, Shimadzu, Columbia, MD) to separate free and conjugated form of miR-145 and quantified for free miR content at λ_ex_ = 650 nm.

### *In vitro* endosomal escape

DSPE-PEG(2000)-Cy5 was incorporated into miR-145 micelles during micelle synthesis at 49:1:40:10 mol ratio of MCP-1 PA, miR-145 amphiphiles, DSPE-PEG(2000)-methoxy, and Cy5 amphiphiles, respectively. hASMC were incubated at a cell density of 3×10^5^ cells/well in Medium 231 (5% SMGS, 1% pen/strep) for 24 hours in 6-well plates containing coverslips. Fluorescently labeled micelles were incubated for 0.5, 1, 2, and 4 hours. The cell suspensions were washed with PBS, fixed with 4% paraformaldehyde, and stained with Lysotracker Green and DAPI, according to manufacturer’s instructions. The cells were observed by a Confocal Laser Scanning Microscope 780 (Zeiss, Oberkochen, Germany) at excitation wavelengths of 405 nm, 504 nm, and 650 nm to visualize the nuclei (blue), acidic compartments of lysosome (green), and micelle internalization (Cy5, red), respectively.

To test endosomal escape, hASMCs were seeded in 24-well plates containing coverslips for 24 hours. Bafilomycin A1 (200 nM, V-ATPase inhibitor) was incubated with hASMCs for 45 minutes prior to incubation of miR-145 micelles or PBS for 4 hours in FBS-free Medium 231. After 4 hours, the cells were replenished with Medium 231 supplemented with SMGS growth medium for an additional 20 hours, prior to PCR analysis of KLF-4, ELK-1, and α-SMA. The co-localization between the endosome/lysosome compartment and Cy5miR-145 micelles were further visualized by confocal microscopy over 0.5, 1, 2, and 4 hours, as described above.

Cy5miR-145 micelles (100 µM) and CT-B Alexa Fluor488 (5 μg/mL) (caveolae-dependent pathway internalization) were simultaneously incubated with VSMCs for 2 hours at 37°C for assessment of caveolin-mediated endocytosis. The cells were stained with DAPI and observed for intracellular fluorescence distribution using confocal microscopy.

### *In vitro* mRNA expression

hASMCs were incubated with nanoparticles or free miRs at a miR concentration of 250 nM in serum-free media. Following the 4 hours transfection period, the media was replaced by supplemented media containing 5% SMGS and incubated for an additional 20 hours. RNA was isolated *via* Trizol (Invitrogen, Carlsbad, CA), and cDNA was synthesized using the RT^2^ First Strand Kit (Qiagen, Hilden, Germany) based on manufacturer’s instructions. Myocardin, α-SMA, calponin, KLF-4/5, ELK-1, and CCR2 expressions were determined by real time-PCR using RT^2^ SYBR Green qPCR Mastermix (Qiagen, Hilden, Germany) on a CFX384 Real-Time PCR Detection System (Bio-Rad Laboratories, Hercules, CA). GAPDH was used as an internal loading control. The 2^−ΔΔC^_T_ method was used to quantify mRNA expression level.

### miR-145 expression and VSMC phenotypic markers on patient-derived VSMCs

The use of human tissues in this study was approved by the Institutional Biosafety Committee (IBC #BUA-16-00057) and Institutional Review Board (IRB #HS-18-00638, BUA-18-0031) at the University of Southern California. Informed consent was obtained from patients for collection of tissue discarded during the normal course of their operation. Segments of the right pulmonary artery and aorta without plaque and from a posterior tibial artery with severe atherosclerosis was collected from male patients. Tissue samples were stored in 1x Hank’s balanced salt solution (HBSS) for no longer than 16 hours prior to VSMC harvest and cell culture. The VSMC harvest and cell culture procedure was adopted from previous reports. Briefly, tissue samples were washed and rinsed three times with HBSS before the endothelium and adventitia were removed, and the medial layer was sectioned into 1-2 mm^2^ explants, digested with 0.25% collagenase type II and 0.5% elastase type II for 16 hours at 37°C. The explant solution was centrifuged at 1500 rpm for 10 min, and the pelleted VSMCs were collected and incubated with 2 mL DMEM supplemented with 10% FBS and 1% penicillin/streptomycin at 37°C for a week or until 70% confluence. VSMCs derived from patients were cultured to passage 3 and were treated with miR-145 micelles and other micelle and miR groups (250 nM miR). miR-145 expression and VSMC phenotypic markers were analyzed by qRT-PCR.

### *In vivo* pharmacokinetics and biodistribution

Male and female adult C57BL/6J wild type mice (8 weeks old) and ApoE^−/−^ (8 weeks old) mice were purchased from The Jackson Laboratory (Bar Harbor, ME). The study protocol was reviewed and approved by the Institutional Animal Care and Use Committee (IACUC) at the University of Southern California. Wild type, eight-week old male (N = 4) C57B1/6J mice (Jackson Laboratory) were intravenously injected with miR-145 micelles containing 10 mol% Cy7 amphiphiles and AF647 miR-145 at a miR dose of 1 mg/kg. Blood was collected at 5 minutes, 1, 3, 8, and 24 hours by orbital sinus, tail vein, or cardiac puncture. Whole blood was centrifuged at 12,000 rpm for 4 minutes to obtain the serum, and miR-145 and amphiphile concentrations were determined by fluorescence measurements (miR-145: excitation: 650 nm; emission: 670 nm and Cy7 amphiphiles: excitation: 730 nm; emission: 783 nm).

At 24 hours, the mice were euthanized and the brain, heart, lung, liver, kidneys, spleen, and bladder were resected and the biodistribution of micelles were quantified by optical imaging (amiHTX, Spectral Instruments Imaging, Tucson, AZ). Signal from the PBS group was used as background measurements.

### *In vivo* evaluation of miR-145 micelles and NT miR-145 micelles biocompatibility and toxicity

Acute toxicity of miR-145 micelles and NT miR-145 micelles was evaluated at the conclusion of the pharmacokinetics studies. Briefly, brain, heart, lung, lung, kidney, spleen, and bladder were harvested and fixed with formalin overnight at 4°C, before submerging in 30% sucrose solution for 8 hours and embedding in OCT at −80°C. OCT-embedded 5 µm tissue sections were stained with H&E and imaged to discern any morphological signs of tissue damage. Blood urea nitrogen (Bioo Scientific, Austin, TX, USA) and urine creatinine (Crystal Chem, Elk Grove Village, IL, USA) levels were measured in mouse serum and urine following manufacturer’s protocols to assess any renal toxicity (N = 4).

### *In vivo* efficacy studies on mid-stage plaque prevention and growth

Eight-week old male (N = 54) or female (N = 54) ApoE^−/−^ mice (The Jackson Laboratory, Bar Harbor, ME) were placed on a Western diet for 14 weeks prior to treatment of miR-145 micelles, miR-67 micelles, NT miR-145 micelles, free miR-145 (1 mg/kg miR (or 0.7 mM total micelle)), or PBS *via* tail vein once every three days for 15 days (5 injections). To check for toxicity, body weight of each mouse was recorded every three days. To check for targeting, micelles were modified to include 10 mol% DSPE-PEG(2000)-Cy7 for the last injection.

### *Ex vivo* aorta imaging and *en face* ORO staining

Mice were euthanized and aortas were perfused with several milliliters of PBS. Aortas were then excised and fluorescently imaged to detect particle fluorescence signal. Next, aortas were fixed in 4% paraformaldehyde overnight at 4°C. Excess adventitial fat on aortas was removed under a stereomicroscope and the tissue was washed in PBS. Aortas were then washed in 78% methanol for 5 minutes under gentle movement for a total for 3 times before incubating in a 2% ORO solution (Sigma-Aldrich, St Louis, MO) for 60 minutes. Aortas were then washed again in 78% methanol twice and cut open longitudinally before imaging.

### *In vivo* prevention of early-stage atherosclerosis

Eight-week old, male (N = 15) or female (N = 15) ApoE^−/−^ mice (Jackson Laboratory) were placed on a Western diet (21% (wt/wt) fat, 0.2% (wt/wt) cholesterol, 19.5% (wt/wt) casein, and no sodium cholate for 4 weeks. After 2 weeks of Western diet, mice were intravenously injected with miR-145 micelles, miR-67 micelles, NT miR-145 micelles, free miR-145 (1 mg/kg miR (or 0.7 mM total micelle)), or PBS *via* tail vein (N = 6). The mice remained on Western diet for two more weeks before euthanasia and evaluating serum concentrations of TNF-α and IL-6 *via* enzyme-linked immunosorbent assay (ELISA, R&D Systems, Minneapolis, MN).

### *In vivo* gene expression

The aortic root region was harvested, weighted out, and lysed with Trizol (Invitrogen, Carlsbad, CA). miR was extracted from aortas of ApoE^−/−^ mice by miRNeasy Mini Kit (Qiagen, Hilden, Germany) and linearly amplified with RT^2^ miR First Strand Kit (Qiagen, Hilden, Germany), according to manufacturer’s instructions. Real-time PCR was conducted on LightCycler 480 Real-Time PCR System (Bio-Rad Laboratories, Hercules, CA) to evaluate cellular miR145, KLF-4 and ELK-1 mRNA levels. GAPDH was used as an internal loading control. The 2^−ΔΔC^_T_ method was used to calculate relative expression changes.

### *In vivo* immunohistochemistry

Hearts, aortic roots, and brachiocephalic arteries were fixed as described above. Per manufacturer’s instructions, serial cross-sections (5 µm) of the aortic root were stained with ORO (Abcam, Cambridge, United Kingdom), H&E, or picrosirius red (Abcam, Cambridge, United Kingdom) for plaque lesion size, necrotic core area, and fibrous cap area, respectively. The area measurements were obtained using ImageJ software. From the observance of aortic valve leaflets to the loss, every 10^th^ aortic root section per animal sample (up to 6 sections) was quantified and averaged to account for differences in plaque structure to obtain a mean ± S.D. measurement.

For fluorescence immunohistochemistry staining of α-SMA, calponin 1, and CD107b, aortic root sections were washed with Tris Buffered Saline (TBS) with 0.025% Triton X-100. The sections were blocked with 1% bovine serum albumin (BSA), 10% normal goat serum, 0.3 M glycine in 0.1% TBS Tween-20 for 1 hour at room temperature, followed by primary antibody staining of diluted anti-alpha smooth muscle actin (Abcam), calponin 1 polyclonal antibody (Bioss Antibodies, Woburn, MA), or rat anti-mouse CD107b (Becton Dickinson Biosciences, Franklin Lakes, NJ) overnight at 4°C in a humidified chamber. The sections were washed and rinsed with TBS, prior to 1 hour incubation with Alexa Fluor 488 anti-mouse IgG (Thermo Fisher Scientific, Waltham, MA) at room temperature in the dark. The samples were counterstained with DAPI and mounted with a coverslip using VectaMount™ aqueous mounting medium (Vector Laboratories, Burlingame, CA).

### Statistical analysis

Results were expressed as means ± standard deviation (S.D.). A two-tailed Student *t*-tests was used to determine statistical significance between two groups, while a one-way analysis of variance (ANOVA) was used to determine statistical significance between more than two groups. A p value < 0.05 was considered statistically significant. All statistical analyses were conducted using GraphPad Prism 8 (GraphPad Software, San Diego, CA).

## General

The authors thank the Center for Electron Microscopy and Microanalysis, Center of Excellence in NanoBiophysics, Core Center of Excellence in Nano Imaging, and Translational Imaging Center at the University of Southern California for assistance in instrumental setups. We also thank our patients for consenting to donate their tissues for this study.

## Funding

This work was support by the University of Southern California, the American Heart Association Predoctoral Fellowship (19PRE34380998) awarded to D.D.C., the National Heart, Lung, and Blood Institute (NHLBI, R00HL124279), New Innovator Award (DP2-DK121328, NIH), Eli and Edythe Broad Innovation Award, the L.K. Whittier Foundation Non-Cancer Translational Research Award, and WISE Gabilan Assistant Professorship granted to E.J.C.

## Author contributions

E.J.C. conceived the project and E.J.C., D.C. and C.P. designed the experiments; C.P., D.C., J.W., J.J., V.O., Z.J., K.C., and E.J.C. performed the experiments and analyzed the data; C.P., D.C, and E.J.C wrote the manuscript. All authors have read and approved the final manuscript.

## Competing interests

There are no competing interests to state.

## Abbreviations

VSMCs: vascular smooth muscle cells
miR-145: micelles miR-145 containing peptide amphiphile micelles
CCR2: C-C chemokine receptor 2
MCP-1: monocyte chemoattractant protein-1
α-SMA: alpha smooth muscle actin
miR: microRNA
KLF-4/5: Krüppel-like factor 4 and 5
ELK-1: ETS domain-containing protein-1
PEG: polyethylene glycol
SRF: serum response factor
TEM: transmission electron microscopy
DLS: dynamic light scattering
CMC: critical micelle concentration
FBS: fetal bovine serum
hASMCs: human aortic smooth muscle cells
Me-β-CD: methyl-β-cyclodextrin
Cy5miR-145: micelles Cy5 labeled-miR-145 micelles
CT-B: cholera toxin subunit B
GSH: glutathione
AF647: Alexa Fluor 647
BUN: blood urea nitrogen
SMGS: smooth muscle growth supplement
IACUC: Institutional Animal Care and Use Committee
HPLC: high performance liquid chromatography
HBSS: Hank’s balanced salt solution
TBS: tris buffered saline
BSA: bovine serum albumin
S.D.: standard deviation
PEI: polyethylenimine
DOTAP: 1,2-dioleoyl-3-trimethylammonium-propane
PCL: polycaprolactone
DOPE: 1,2-dioleoyl-sn-glycero-3-phosphoethanolamine
PLL: poly(L-lysine)
DMTAP: dimyristoyltrimethylammonium propane
DMPC: dimyristoylphosphatidylcholine
BHEM/chol: N,N-bis(2-hydroxyethyl)-N-methyl-N-(2-cholesteryloxycarbonyl aminoethyl) ammonium bromide.

## Supplementary Materials

**Fig. S1.**
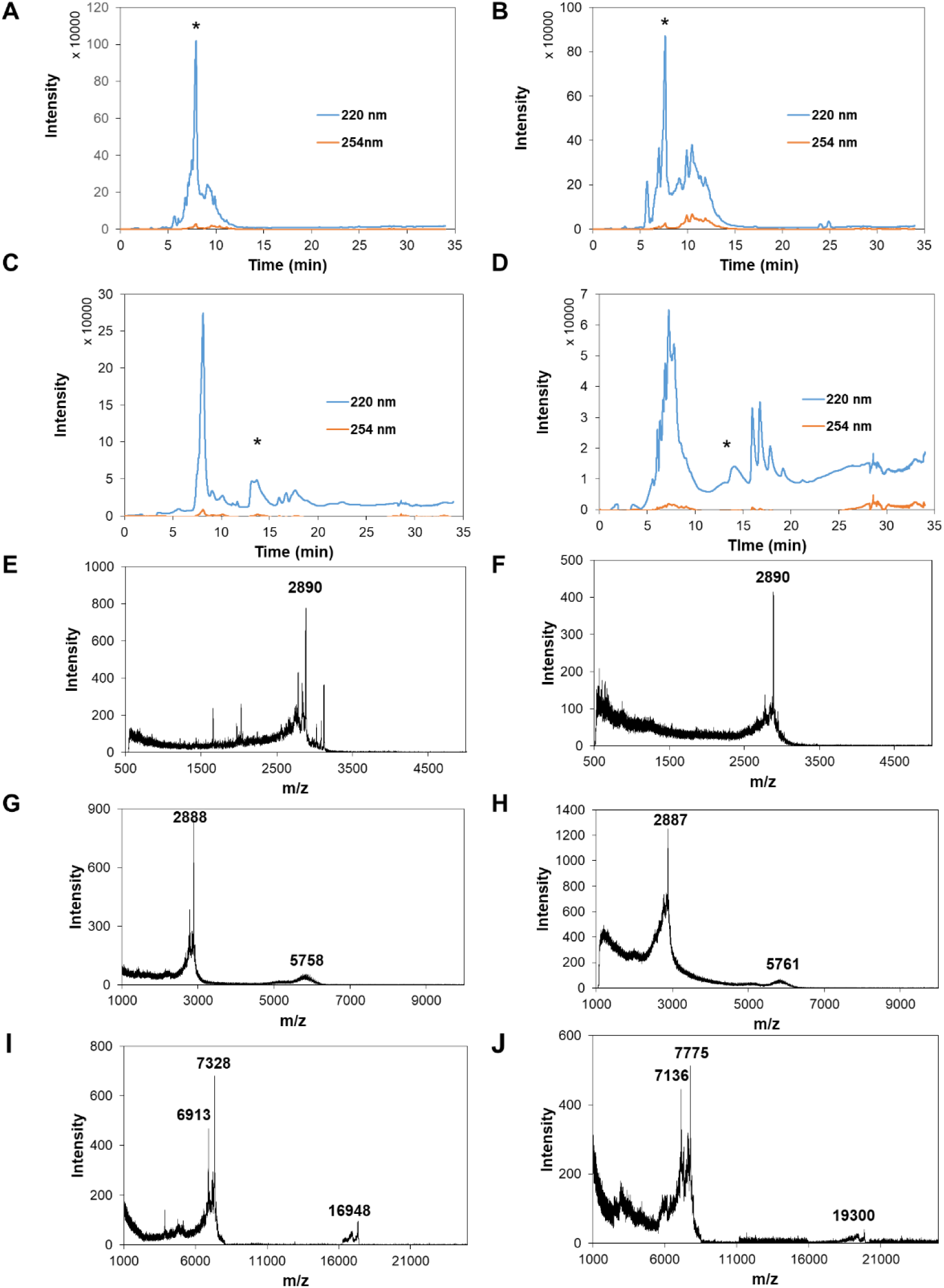
HPLC chromatograms of (A) MCP-1, (B) NT peptides, (C) DSPE-PEG(2000)-MCP-1 and (D) DSPE-PEG(2000)-N at 220 nm (average peptide bond wavelength) and 254 nm (tyrosine wavelength). Highlighted peaks at 7.4 min and 7.2 min signified the elution of the MCP-1 and NT peptides, respectively, and peaks at 16.9 min signified the elution of the DSPE-PEG(2000)-MCP-1 and MCP-1 and DSPE-PEG(2000)-NT. MALDI-TOF mass spectra of (E) MCP-1 and (F) NT peptides confirmed the correct molecular weight of 2890 Da, and (G) DSPE-PEG(2000)-MCP-1 and (H) DSPE-PEG(2000)-NT of the correct molecular weights of 5761 Da. MALDI-TOF mass spectrum of (I) DSPE-PEG(2000)-miR-145 and (J) DSPE-PEG(2000)-miR-67 at a range of 1000-25000 Da showing the peaks for anti-sense strand at m/z 6913 and 7136, sense strand at m/z 7328 and 7775, and DSPE-PEG(2000)-miR at m/z 16948 and 19300 (Expected: 17047 Da and 17717 Da).

**Fig. S2.**
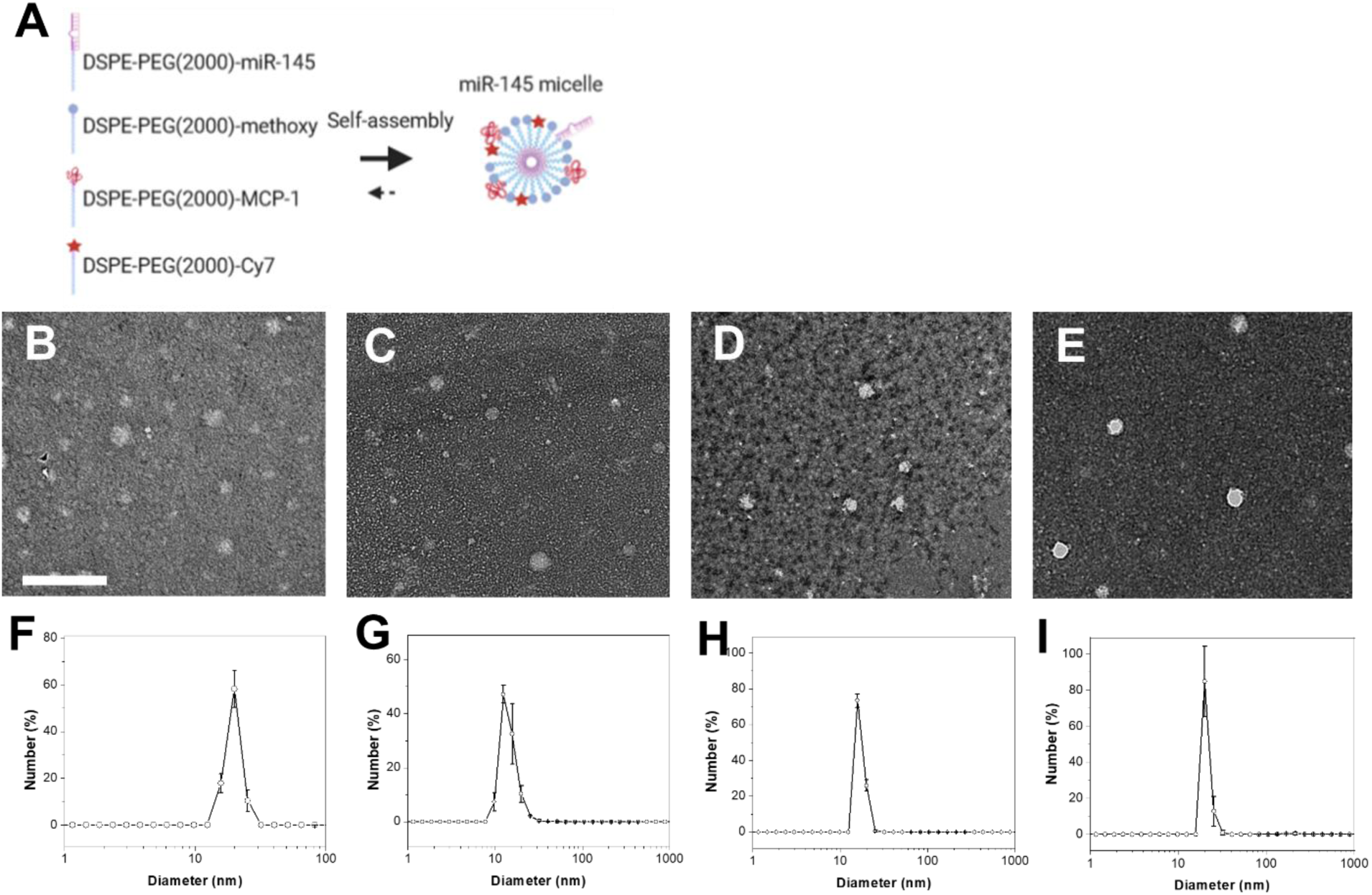
Preparation and characterization of miR-145 micelles. (A) Schematic representation of the self-assembly and multifunctionality of miR-145 micelles. (B, C, D, E) TEM images of miR-145 micelles, MCP-1 micelles, miR-67 micelles, and NT miR-145 micelles, respectively. Scale bar = 200 nm. (F, G, H, I) Number-average size distribution of miR-145 micelles, MCP-1 micelles, miR-67 micelles, and NT miR-145 micelles, respectively.

**Fig. S3.**
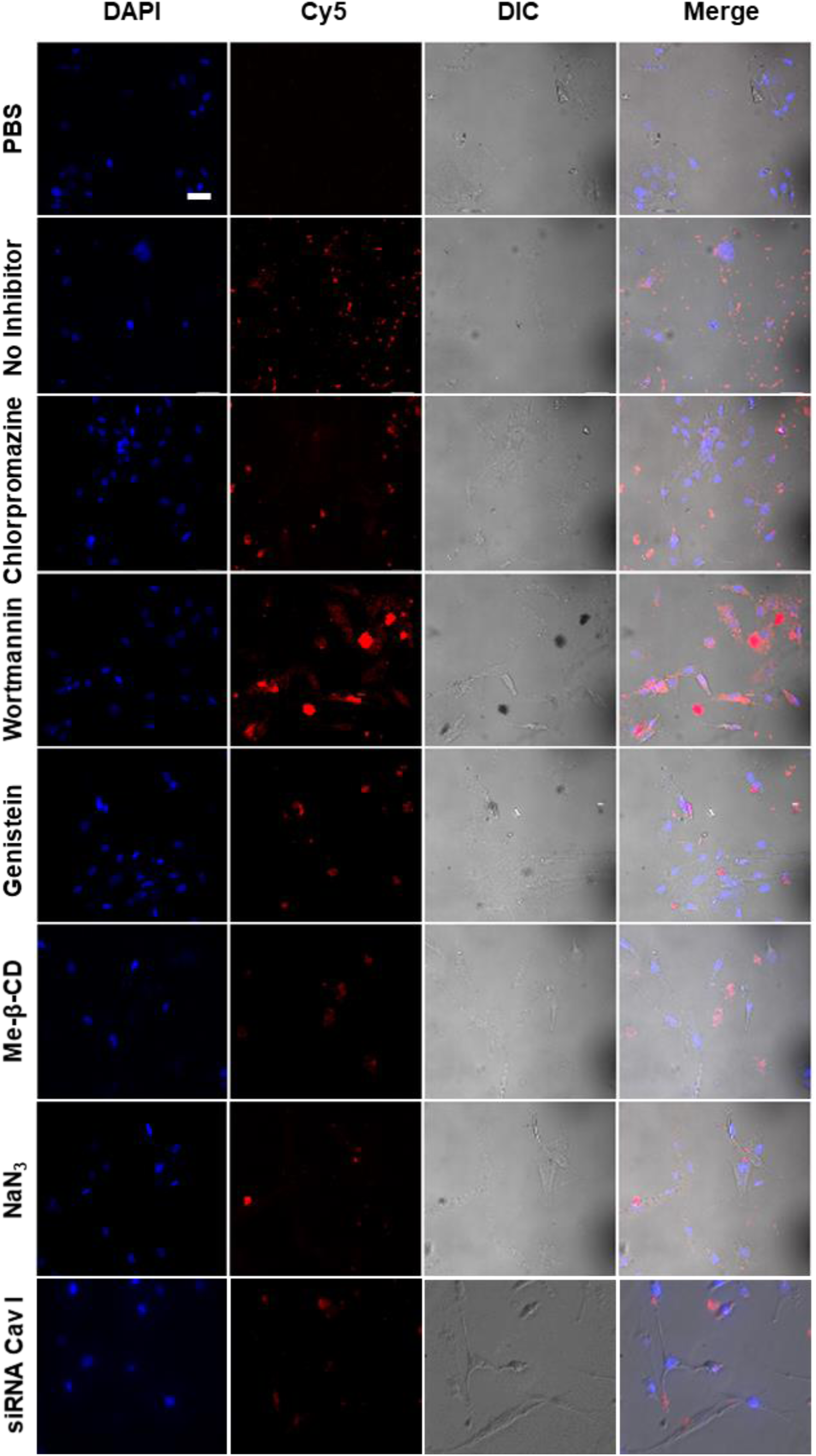
Fluorescence images of hASMCs incubated with miR-145 micelles with different endocytosis inhibitors: chlorpromazine (clathrin-mediated endocytosis), wortmannin (transferrin receptor endocytosis), genistein (caveolae-mediated endocytosis), me-β-CD (lipid raft-mediated endocytosis), sodium azide (energy-dependent endocytosis), and siRNA caveolin I. Channels: Cy5 micelles (red) and DAPI (blue). Scale bar = 50 µm.

**Fig. S4.**
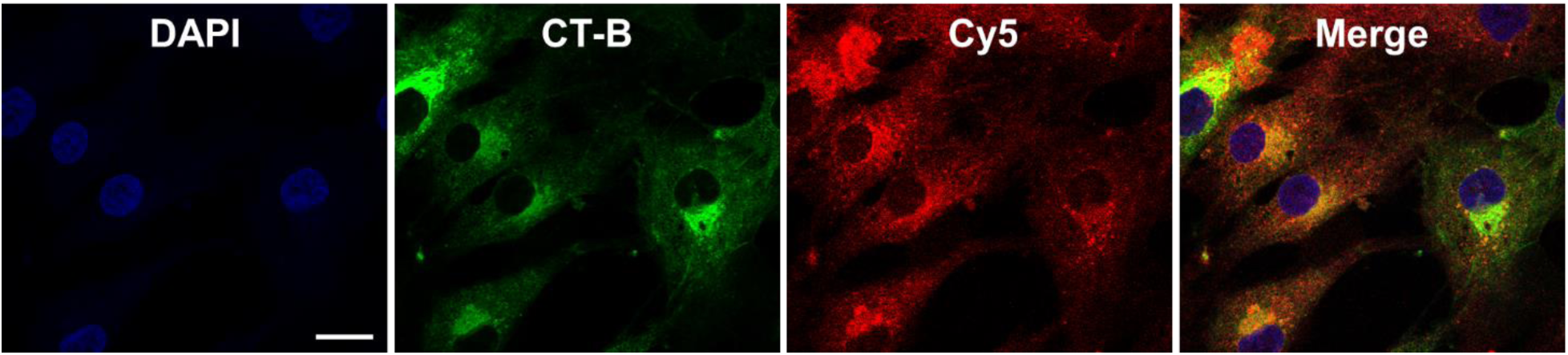
Fluorescence images of hASMC incubated with miR-145 micelles and treated with CT-B confirming micelle internalization within hASMCs is through caveolae-mediated endocytosis. Channels are: Cy5-micelles (red) and CT-B (green, known for internalization *via* caveolae-mediated endocytosis). Scale bars = 20 μm.

**Fig. S5.**
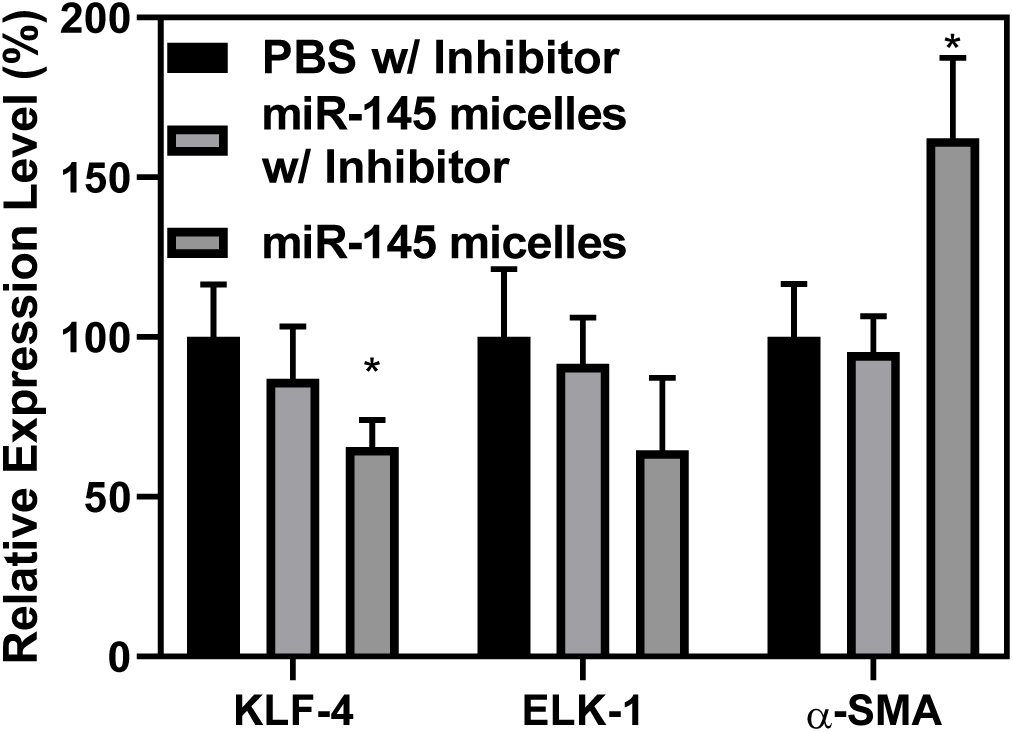
mRNA expression of KLF-4, ELK-1, and α-SMA in hASMC transfected with miR-145 micelle (miR concentration of 250 nM) and inhibited with bafilomycin A1 demonstrating proton sponge effect plays a role in the endosomal escape of miR-145 micelles (N = 3; *p < 0.05).

**Fig. S6.**
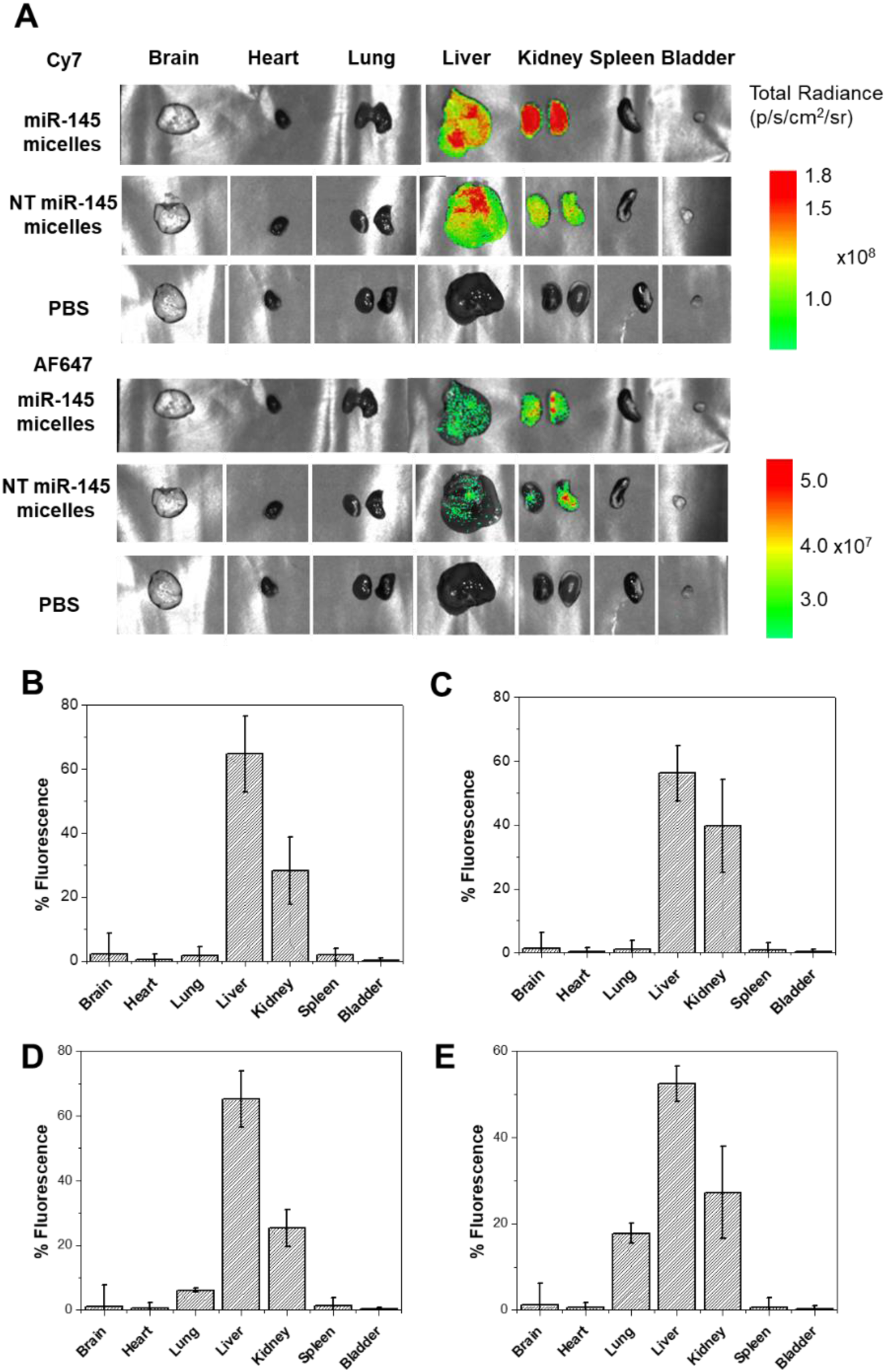
(A) Tissue biodistribution of Cy7 and AF647 determined by optical imaging of miR-145 micelle, NT-miR-145 micelle, and AF647 labeled miRs in C57BL/6J mice 24 hours post injection. (B) Cy7 and (C) AF647 distribution in tissues 24 hours after intravenous administration of miR-145 micelle or (D) Cy7 and (E) AF647 distribution NT-miR-145 micelle (N = 4).

**Fig. S7.**
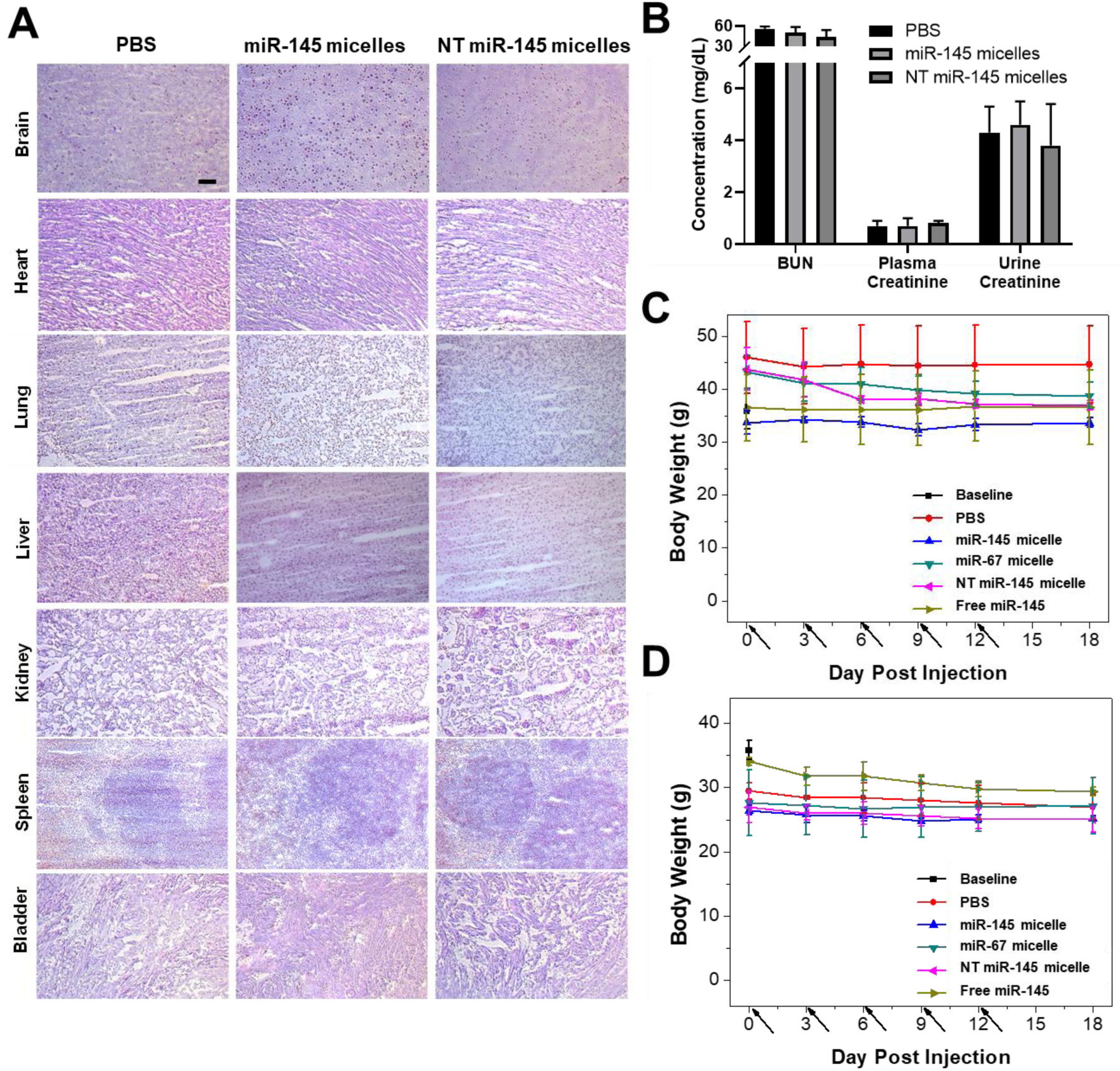
(A) H&E stained images of organs from C57BL/6J mice treated with PBS, miR-145 micelles, or NT-miR-145 micelles showing micelles are non-toxic *in vivo*. Scale bar = 100 µm. (B) BUN and creatinine measurements in PBS, miR-145 micelle, and NT miR-145 micelle treated C57BL/6J mice (N = 4). Body weight evolution in (C) male and (D) female mid-stage atherosclerotic mice treated with PBS (●), miR-145 micelles (▲), miR-67 micelles (▼), NT miR-145 micelles (◄), or miR-145 (►) at an miR dose of 0.3 mg /kg on day 0, 3, 6, 9, and 12 (N = 6). No significant change in body weight was observed for micelle-treated groups, demonstrating the safety and tolerance of micelles.

**Fig. S8.**
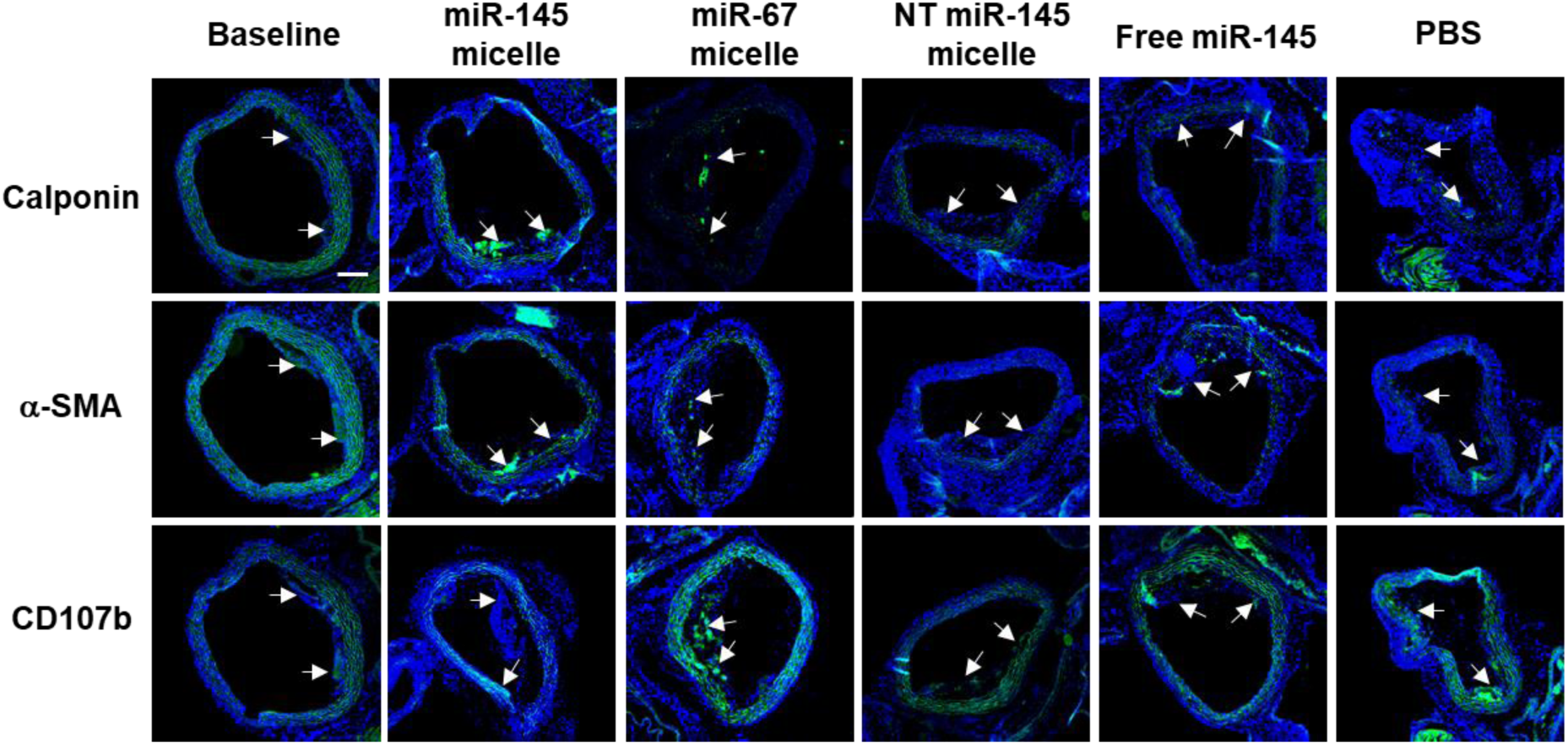
Immunohistochemistry staining using calponin, α-SMA, and CD107b antibodies of the aorta in ApoE^−/−^ mice treated with miR-145 micelles, miR-67 micelles, NT miR-145 micelles, free miR-145, or PBS in mid-stage atherosclerotic mice. miR-145 micelles showed an increase in calponin and α-SMA contractile markers and a decrease in CD107b+ macrophages within atherosclerotic lesions. Scale bar = 200 µm.

**Fig. S9.**
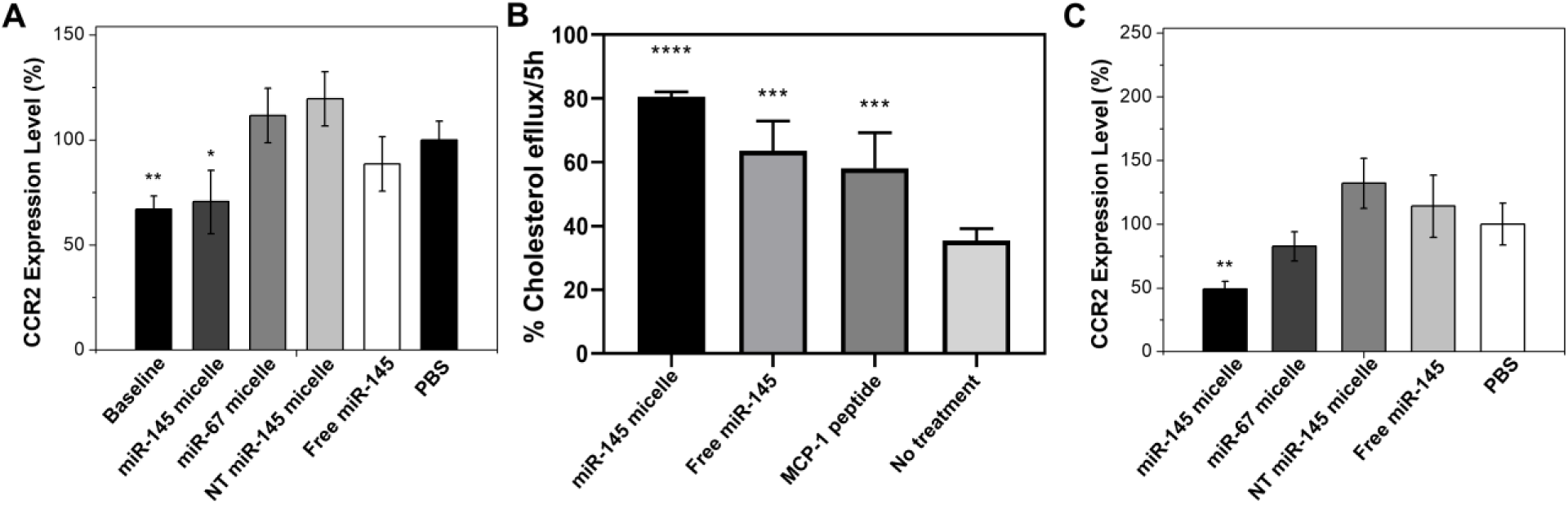
miR-145 micelles decrease CCR2 expression within the aortic root in the (A) mid-stage atherosclerosis model (N = 6; *p < 0.05, **p < 0.01). (B) *In vitro* cholesterol efflux in hASMCs after cholesterol activation for 48 hours and treatment with miR-145 micelles, free miR-145 or MCP-1 peptide highlighting the cholesterol efflux capabilities of miR-145 micelles (N = 6). miR-145 micelles decrease CCR2 expression within the aortic root in the (C) early-stage atherosclerosis model (N = 6; **p < 0.01).

**Fig. S10.**
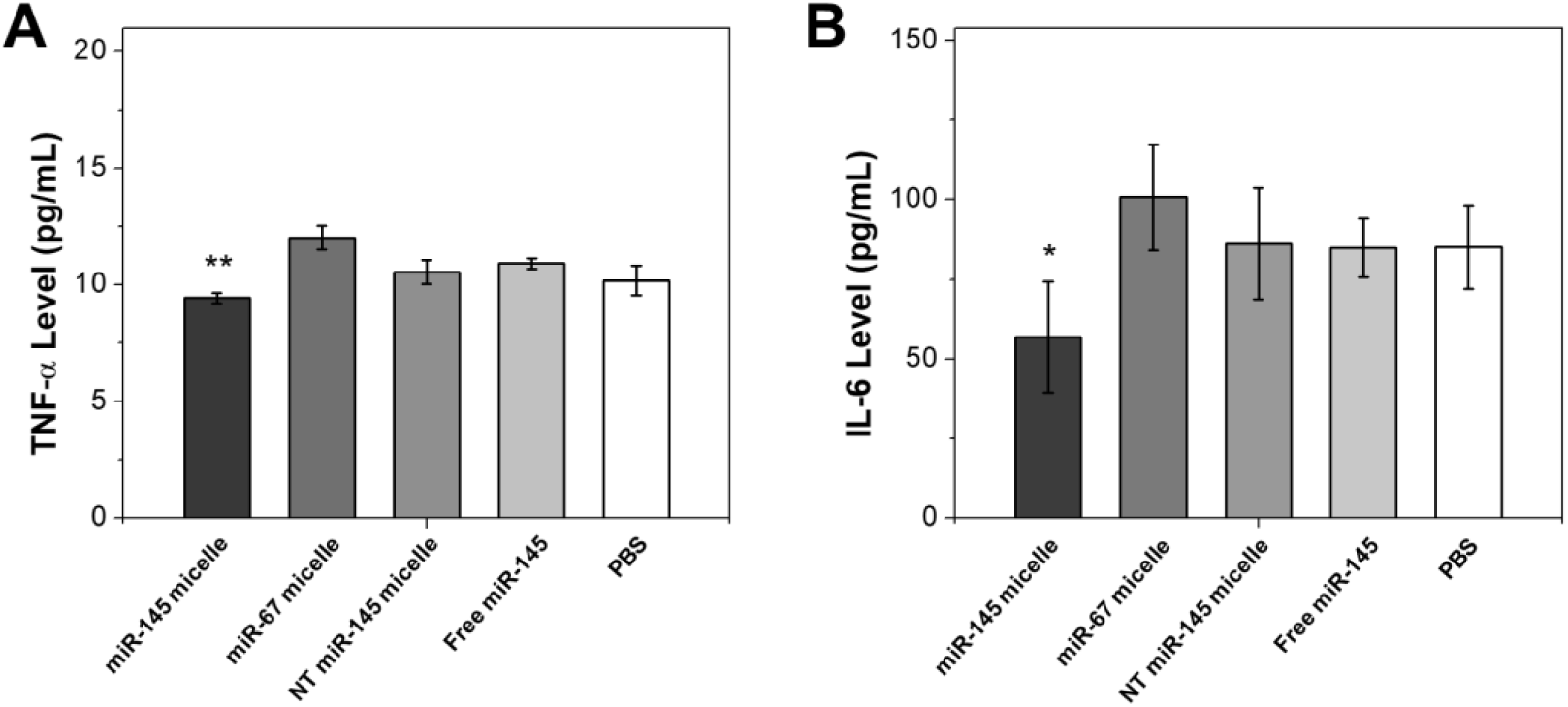
Mice treated with miR-145 micelles show a decrease in inflammatory and immunogenic responses. (A) TNF-α and (B) IL-6 atherogenesis promoting cytokine levels in serum of mice treated with miR-145 micelles, miR-67 micelles, NT miR-145 micelles, miR-145, or PBS in early-stage preventive studies (N = 6; *p < 0.05, **p < 0.01).

## Notes

### Competing Interest Statement

The authors have declared no competing interest.

